# Post-transcriptional regulation by the CCR4-NOT deadenylase complex maintains redox homeostasis in insulin biosynthesis in mouse pancreatic β cells

**DOI:** 10.1101/2024.06.26.599433

**Authors:** Akiko Yanagiya, Patrick N. Stoney, Yuki Tara, Shohei Takaoka, Guillaume Vares, Rei Higa, Nicholas R. Friedman, Alejandro Villar-Briones, Tadashi Yamamoto

## Abstract

Pancreatic β cells synthesize insulin to maintain glucose homeostasis. In diabetes, elevated blood glucose and insulin resistance compel β cells to produce insulin, and hence β cells are vulnerable to oxidative stress by glucotoxicity. In insulin biosynthesis, the conversion of proinsulin to insulin is initiated by forming disulfide bonds in proinsulin for oxidative protein folding. Insulin content and insulin secretion in β cells are decreased by deletion of CNOT7, a catalytic subunit of the CCR4-NOT complex, and accompanied by increased proinsulin, implying the impaired conversion of proinsulin to insulin. We found that PRDX4 essential for disulfide bond formation in proinsulin, is reduced in *Cnot7*-KO β cells. Moreover, protein expression of CNOT8, a paralog of CNOT7, is increased in *Cnot7*-KO β cells and binds to *Prdx4* mRNA via MSI2. Here, we demonstrate the post-transcriptional regulation of *Prdx4* mRNA by the CCR4-NOT complex to maintain oxidative-reductive homeostasis in insulin biosynthesis.

**HIGHLIGHTS:**

- Insulin content and secretion are decreased in *Cnot7*-KO pancreatic β cells.
- PRDX4, needed for proinsulin folding, is decreased in *Cnot7*-KO pancreatic islets.
- The conversion of proinsulin to insulin is impaired in *Cnot7*-KO β cells.
- CNOT8, but not CNOT7, interacts with MSI2 that binds to *Prdx4* mRNA

## INTRODUCTION

Post-transcriptional regulation plays pivotal roles in a variety of physiological processes because it allows a rapid response to external and/or internal environmental changes to maintain cell homeostasis without *de novo* RNA synthesis. The CCR4-NOT deadenylase complex serves as a 3’-5’ exonuclease by removing the poly(A) tail at the 3’ end of mRNAs at the first stage of mRNA degradation (Shirai et al., 2014). Our past studies have revealed its importance in metabolic functions such as resistance to obesity (Morita et al., 2011) (Takahashi et al., 2015) and pancreatic β cell identity (Mostafa et al., 2020b).

The catalytic subunits of the CCR4-NOT complex, CNOT7 and CNOT8, have been thought to be redundant because of their highly similar amino acid sequences and similar characteristics in *in vitro* experiments (Aslam et al., 2009). CNOT7 and CNOT8 might possess distinctive physiological functions in embryogenesis since *Cnot8*-whole-body knockout (*Cnot8*-KO) mice are embryonic lethal whereas *Cnot7*-KO mice are viable (Nakamura et al., 2004) (Mostafa et al., 2020a). This different requirement for CNOT7 and CNOT8 in embryogenesis prompted us to investigate whether CNOT7 and CNOT8 might have distinct roles in other physiological functions. Although *Cnot7*-KO mice are resistance to age- and diet-induced obesity with enhanced insulin sensitivity (Takahashi et al., 2015), little is known about the function of CNOT7 in insulin biosynthesis, which is a key biological process to maintain blood glucose homeostasis. Notably, pancreatic β cell-specific disruption of *Cnot3* (*Cnot3*-βKO), encoding a regulatory subunit of the CCR4-NOT complex, results in diabetes with hyperglycemia and impaired insulin secretion due to loss of pancreatic β cell identity (Mostafa et al., 2020b). Allowed genes and disallowed genes that are required to be increased and decreased, respectively, in the process of pancreatic β cell maturation were dysregulated in *Cnot3*-βKO pancreatic islets.

Insulin is a hormone that maintains blood glucose homeostasis by facilitating glucose uptake into cells. Insulin is solely synthesized and secreted by pancreatic β cells, which are highly specialized for insulin production. Translation of *insulin* mRNA generates a single polypeptide called preproinsulin in the cytoplasm. Newly synthesized preproinsulin is transferred to the endoplasmic reticulum (ER) by a signal peptide at the N-terminus, where the signal peptide is removed to generate proinsulin (Dodson and Steiner, 1998) (Liu et al., 2014) (Liu et al., 2018). Relative to mature insulin, proinsulin possesses low biological activity to reduce blood glucose. The conversion of proinsulin to insulin is initiated by oxidative folding of proinsulin to form a three-dimensional structure stabilized by three disulfide bonds, followed by a removal of a short peptide (the C-peptide) by prohormone convertase 1/3 (PCSK1), prohormone convertase 2 (PCSK2) and carboxypeptidase E (CPE). Disulfide bond formation is crucial for the proper folding of proinsulin and the subsequent biological action of insulin to reduce blood glucose levels. This oxidative folding is accomplished by protein disulfide isomerase (PDI), endoplasmic reticulum oxidase 1 (ERO1) and peroxiredoxin-4 (PRDX4). Hyperglycemia and insulin resistance caused by diabetes chronically stimulate pancreatic β cells to produce and secrete insulin in an attempt to rectify blood glucose homeostasis, eventually overwhelming antioxidative and protein-folding capacities and impairing ER redox homeostasis in β cells (Laybutt et al., 2007) (Fonseca et al., 2009). PRDX4 is an ER-specific antioxidative peroxidase that oxidizes PDI and promotes oxidative folding of proinsulin, thereby contributing to disulfide bond formation in proinsulin. PRDX4 also positively regulates insulin biosynthesis and glucose-induced insulin secretion in pancreatic β-cells and insulin-producing cells (Ding et al., 2010) (Mehmeti et al., 2014) (Tran et al., 2020).

Here, we deciphered the molecular mechanism of the post-transcriptional regulation of *Prdx4* mRNA by the CCR4-NOT complex containing CNOT8, but not CNOT7, to maintain redox homeostasis to form disulfide bonds in proinsulin, contributing to functional insulin biosynthesis in pancreatic β cells.

## RESULTS

### *Cnot7*-KO mice exhibit impaired insulin secretion

To elucidate the metabolic phenotypes of 8-week-old *Cnot7*-KO mice, intraperitoneal glucose tolerance test (IPGTT) and insulin tolerance test (ITT) were conducted. There was a slight difference in glucose tolerance between wild-type (WT) and *Cnot7*-KO mice (Figures 1A and 1B), whereas insulin sensitivity was significantly enhanced in *Cnot7*-KO mice compared to WT mice (Figures 1C and 1D). Moreover, there was no difference in body weight, fed blood glucose and fasting blood glucose between WT and *Cnot7*-KO mice (Figures S1A, S1B and S1C). Notably, serum insulin was reduced in *Cnot7*-KO mice (Figure 1E) in accordance with impaired glucose-stimulated insulin secretion (GSIS) in *Cnot7*-KO mice (Figure 1F). Immunostaining of pancreas sections suggested that insulin is reduced in *Cnot7*-KO β cells (Figure 1G), but there was no difference between WT and *Cnot7*-KO mice in terms of pancreatic weight (Figure S1D), the morphology of islets (Figure S1E), islet area (Figure S1F), the morphology of pancreatic β cells (Figure S1G) and β cell area (Figure S1H). In accordance with morphological observations, the β cell mass of *Cnot7*-KO mice was similar to that of WT mice (Figure 1H), whereas insulin content and GSIS using isolated islets were significantly reduced in *Cnot7*-KO mice (Figures 1I and 1J). These data suggest that *Cnot7*-KO mice have a specific defect in insulin content and secretion in pancreatic β cells.

**Figure 1.**
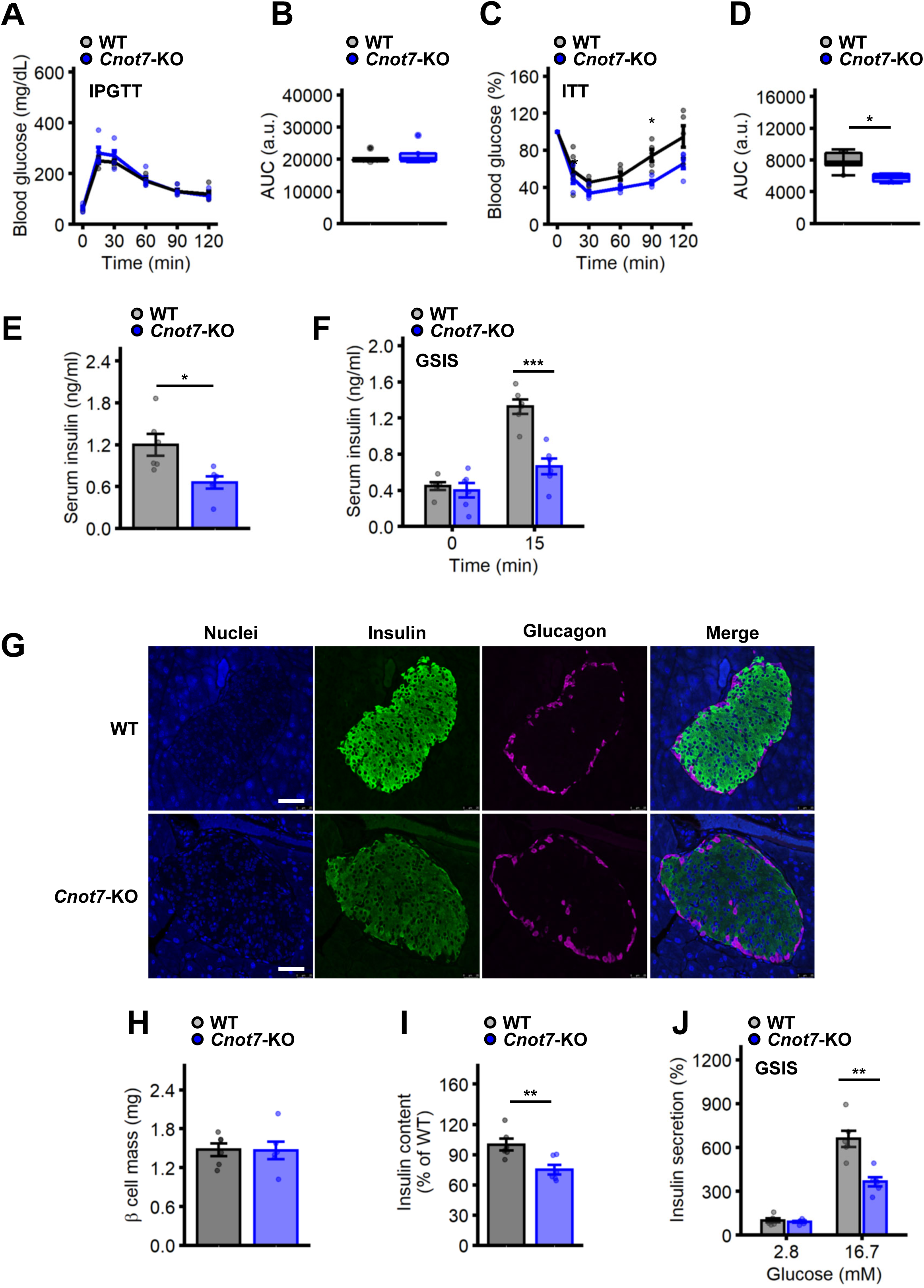
Insulin secretion from pancreatic β cells of *Cnot7*-KO mice is impaired. **(A)** Intraperitoneal glucose tolerance test (IPGTT) using 8-week-old WT (gray, n=5) and *Cnot7*-KO (blue, n=5) mice. Blood glucose levels (mg/mL) were measured at 0, 30, 60, 90 and 120 minutes after administration of 1g glucose per kg of body weight. **(B)** Areas under the curves (AUC) of IPGTT in Figure 1A are shown. **(C)** Insulin tolerance test (ITT) using 8-week-old WT (gray, n=5) and *Cnot7*-KO (blue, n=5) mice. Blood glucose levels were measured at 0, 30, 60, 90 and 120 minutes after insulin intraperitoneal administration (1U insulin per kg of body weight). Values are shown as a percentage of the value at 0 minute. **(D)** AUCs of ITT in Figure 1C are shown. **(E)** Serum insulin of fed 8-week-old WT (gray, n=5) and *Cnot7*-KO (blue, n=5) mice. **(F)** Glucose-stimulated insulin secretion (GSIS) in 8-week-old WT (gray, n=5) and *Cnot7*-KO (blue, n=5) mice. After mice were fasted for 16 hours, serum insulin levels (ng/ml) were measured at 0 and 15 minutes after glucose administration. **(G)** Immunofluorescent labeling of insulin (green) and glucagon (magenta) in islets of 8-week-old WT and *Cnot7-*KO mice. Nuclei were labelled with DAPI (blue). Scale bar = 50 µm. **(H)** Pancreatic β cell mass (mg) of 8-week-old WT (gray, n=5) and *Cnot7-*KO (blue, n=5). **(I)** Insulin content (% of WT) of 8-week-old WT (gray, n=5) and *Cnot7-*KO (blue, n=5) mice. **(J)** GSIS in isolated islets of 8-week-old WT (gray, n=5) and *Cnot7-*KO (blue, n=5) mice. Isolated islets were incubated either low (2.8 mM) or high (16.7 mM) glucose for 1 hour. Data represent means ±SEM. Two-way ANOVA followed by Bonferroni post hoc test for IPGTT and ITT. Unpaired Student’s t-test for AUC and other experiments, *p<0.05; **p<0.01; and ***p<0.001.

### *Prdx4* mRNA is decreased in *Cnot7*-KO pancreatic islets

To identify differentially expressed genes in *Cnot7*-KO islets compared to WT, polyadenylated RNAs isolated from islets were analyzed by RNA-seq. Significantly up-regulated (158) and down-regulated (199) transcripts were identified by RNA-seq (Figures 2A, 2B and 2C). Gene ontology (GO) using significantly down-regulated transcripts revealed that mRNAs encoding proteins related to metabolism are enriched in the down-regulated genes (Figure 2D). Intriguingly, the oxidation-reduction process, which is essential for the formation of disulfide bonds during the crucial step of proinsulin folding, was enriched. In addition to RNA-seq analysis, essential transcripts for insulin biosynthesis such as *Ins1*, *Ins2*, *Pcsk1*, *Pcsk2* and *Cpe* were analyzed by qPCR (Figure 2E) in WT and *Cnot7*-KO islets, revealing that these transcripts were not altered. This suggests that reduced insulin content and impaired insulin secretion in *Cnot7*-KO islets are not caused by the reduction of these transcripts. Next, expression of transcripts needed for oxidation-reduction processes such as *Pdi*, *Ero1*, *Ero1b* and *Prdx4* was assessed by qPCR (Figure 2F). Relative mRNA expression of *Pdi*, *Ero1* and *Prdx4* was significantly reduced, implying that the redox system is altered in *Cnot7*-KO islets. In addition, relative mRNA expression of components of the CCR4-NOT complex was evaluated by qPCR (Figure S2B). As expected, relative *Cnot7* mRNA expression was decreased in *Cnot7*-KO islets whereas expression of other components was not altered. mRNA expression of allowed genes that are expressed in mature β cells such as *Mafa* and *Pdx1* was similar between WT and *Cnot7*-KO islets, suggesting that loss of CNOT7 does not affect β cell maturation (Figure 2G) (Kaneto et al., 2008). Moreover, mRNA expression of disallowed genes that are suppressed in mature β cells such as *Ldha*, *Aldob*, *Acot7*, *Hk1*, *Slc16a1* and *Wnt5b* was not increased, revealing that the maturation of β cells was not impaired by the lack of CNOT7 (Figure 2H) (Martinez-Sanchez et al., 2016; Mostafa et al., 2020a; Pullen et al., 2010) (Dhawan et al., 2015).

**Figure 2.**
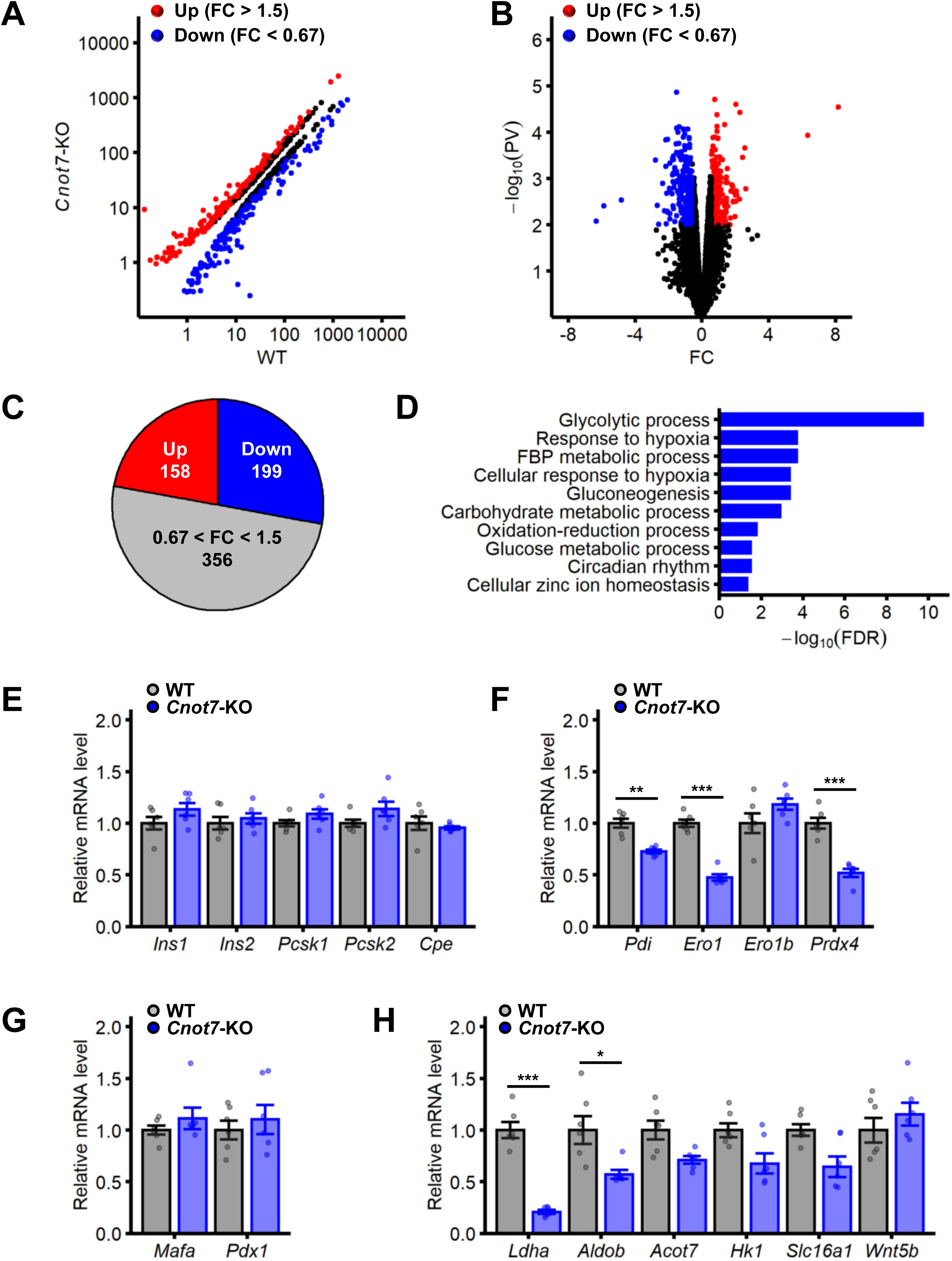
mRNAs encoding genes that are essential for a redox system are down-regulated in *Cnot7*-KO islets. **(A)** A scatter plot of the differentially expressed transcripts (p < 0.01) identified by RNA-seq. between WT (n=3) and *Cnot7*-KO (n=3) islets. Significantly up-regulated transcripts (red; fold change (FC) > 1.5) and down-regulated ones (blue; FC < 0.67) are shown. **(B)** A volcano plot of the differentially expressed transcripts (p < 0.01) between WT and *Cnot7*-KO islets determined by RNA-seq. Significantly up-regulated transcripts (red; fold change (FC) > 1.5) and down-regulated ones (blue; FC < 0.67) are shown. **(C)** A pie chart of transcripts of significantly up-regulated (FC > 1.5; red; 158 transcripts), down-regulated (FC < 0.67; blue; 199 transcripts) and non-significant (0.67 < FC < 1.5; gray; 356 transcripts) in *Cnot7*-KO (n=3) islet relative to WT (n=3), identified by RNA-seq. **(D)** Gene ontology (GO) enrichment analysis (false discovery rate, FDR < 0.05) for down-regulated transcripts. A bar plot of GO terms of biological process ranked by FDR is shown. FBP, fructose 1,6-bisphosphate. **(E-H)** Relative expression of mRNAs that are essential for insulin biosynthesis (*Ins1*, *Ins2*, *Pcsk1*, *Pcsk2* and *Cpe*) **(E)**, oxidation-reduction (*Pdi*, *Ero1*, *Ero1b* and *Prdx4*) **(F)**, allowed genes (*Mafa* and *Pdx1*) **(G)** and disallowed genes (*Ldha*, *Aldob*, *Acot7*, *Hk1*, *Slc16a1* and *Wnt5b*) **(H)** between WT (gray; n=6) and *Cnot7*-KO (blue; n=6) islets, determined by quantitative PCR (qPCR). Average values of WT were set as 1. Data represent means ±SEM. Unpaired Student’s t-test, *p<0.05; **p<0.01; and ***p<0.001.

We assumed that CNOT7 may influence mRNA expression of *Pdi*, *Ero1* and *Prdx4* via both transcription and degradation. Pre-mRNA expression of *Ins1*, *Ins2*, *Pcsk2*, *Pdi* and *Prdx4*, determined by qPCR, was not affected by loss of CNOT7 (Figure S2C), suggesting that altered mRNA expression was likely to result from changes in mRNA stability. To investigate the stability of *Ins1*, *Pcsk2*, *Pdi* and *Prdx4* mRNAs, WT and *Cnot7*-KO islets were treated with a transcriptional inhibitor, actinomycin D, for 0, 4 and 8 hours, and their relative mRNA expression was evaluated by qPCR. *Pdi* and *Prdx4* mRNAs were less stable in *Cnot7*-KO islets than WT, whereas *Ins1* and *Pcsk2* mRNAs were not different (Figures 3A, 3B, 3C and 3D). Stability of *Ins2*, *Pcsk1*, *Cpe* and *Ero1* was also determined by qPCR, revealing that *Ero1* mRNA is destabilized by the lack of CNOT7 (Figures S3A, S3B, S3C and S3D). These data suggest that stability of mRNAs encoding proteins regulating oxidation-reduction processes are reduced by loss of CNOT7.

**Figure 3.**
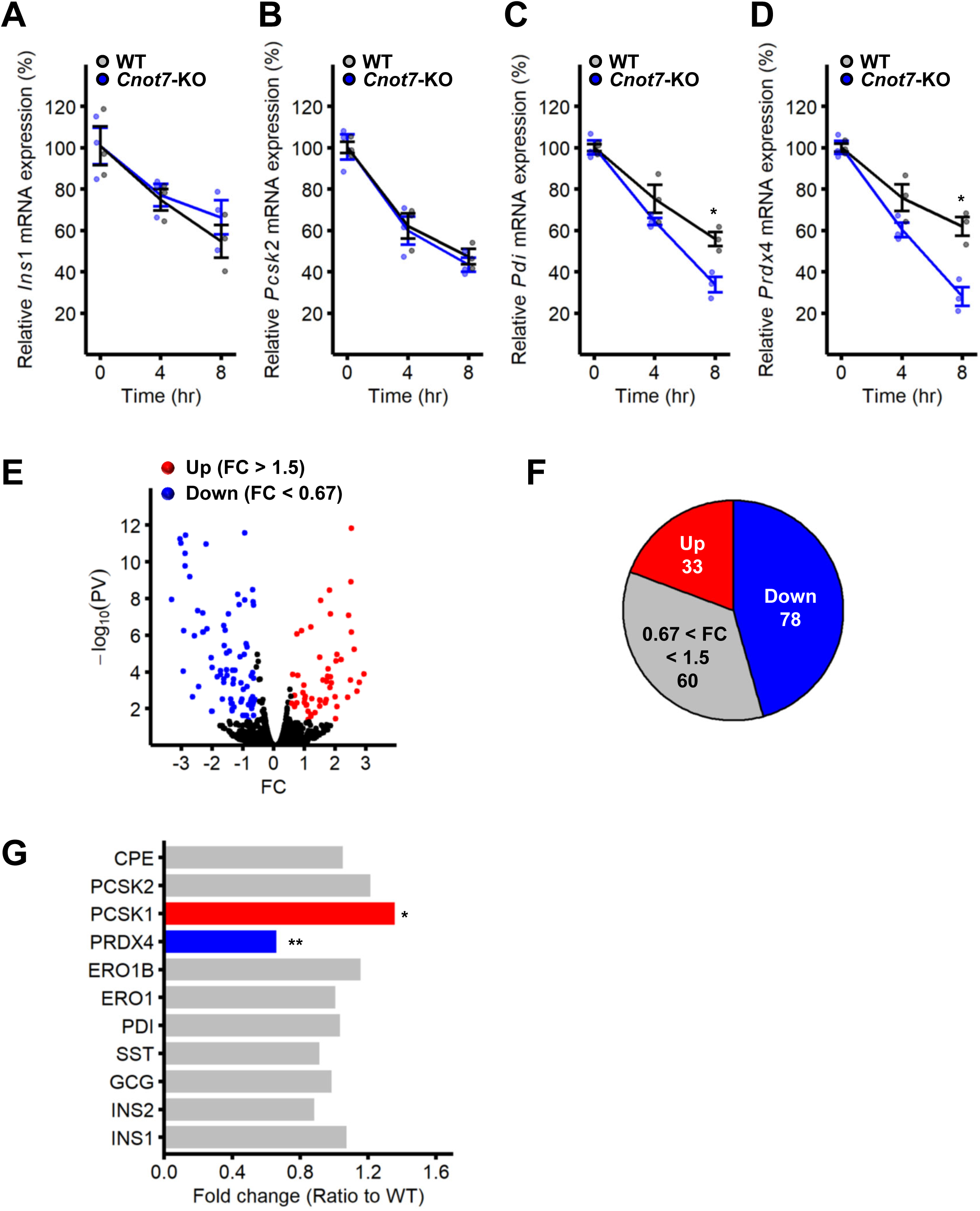
PRDX4 is down-regulated in *Cnot7*-KO islets. **(A-D)** mRNA stability of *Ins1* **(A)**, *Pcsk2* **(B)**, *Pdi* **(C)** and *Prdx4* **(D)** was determined by qPCR using islets of 8-week-old WT (gray; n=3) and *Cnot7*-KO (blue; n=3) mice treated with actinomycin D for 0, 4 and 8 hours. Average values of WT mice at 0 hour were set as 100%. Data represent means ±SEM. Two-way ANOVA followed by Bonferroni post hoc test, *p<0.05; **p<0.01; and ***p<0.001. **(E)** A volcano plot of the differentially expressed proteins (p < 0.05) between WT and *Cnot7*-KO islets determined by mass spectrometry. Significantly up-regulated proteins (red; fold change (FC) > 1.5) and down-regulated ones (blue; FC < 0.67) are shown. **(F)** A pie chart of significantly up-regulated (FC > 1.5; red; 33), down-regulated (FC < 0.67; blue; 78) and non-significant (0.67 < FC < 1.5; gray; 60) proteins identified by mass spectrometry between WT (n=3) and *Cnot7*-KO (n=3) islets. **(G)** A bar plot of protein expression determined by mass spectrometry between WT (n=3) and *Cnot7*-KO (n=3) islets. Significantly up-regulated PCSK1 (red) and down-regulated PRDX4 (blue) are indicated. Data represent means ±SEM. Unpaired Student’s t-test, *p<0.05; **p<0.01; and ***p<0.001.

### PRDX4 is reduced in *Cnot7*-KO pancreatic islets

Differentially expressed proteins in *Cnot7*-KO islets compared to WT were analyzed by mass spectrometry (MS). Significantly up-regulated (33) and down-regulated (78) proteins were identified (Figures 3E and 3F). Notably, PRDX4 was decreased whereas other key proteins that are essential for insulin biosynthesis were not decreased by the lack of CNOT7 (Figure 3G), indicating the potential role of PRDX4 in the phenotype in *Cnot7*-KO mice. Since PRDX4 is reduced at both mRNA and protein levels, further experiments focus on the molecular mechanism by which decreased PRDX4 in *Cnot7*-KO β cells results in impaired insulin biosynthesis. The data suggested that ER redox homeostasis, which is essential for insulin biosynthesis, might be dysregulated by decreased PRDX4 in *Cnot7*-KO β cells.

### Conversion of proinsulin to insulin is impaired in *Cnot7*-KO pancreatic β cells

Since PRDX4 is known to interact with proinsulin and contributes to the formation of disulfide bonds inside proinsulin, the conversion of proinsulin to insulin in *Cnot7*-KO islets was evaluated by western blotting (Figure 4A). Notably, CNOT8, a paralog of CNOT7, was increased in *Cnot7*-KO islets whereas another component of the CCR4-NOT complex, CNOT3, was not altered (Figures 4B). In accordance with MS data, PRDX4 was reduced in *Cnot7*-KO islets (Figure 4C), and this is likely to contribute to the impaired conversion of proinsulin to insulin in the absence of CNOT7 (Figure 4D). *Ins1*, *Ins2* and *Cnot8* mRNAs were not reduced in *Cnot7*-KO islets, suggesting that both impaired conversion of proinsulin and increased CNOT8 are caused by post-transcriptional regulation (Figure 4E). It is feasible that impaired conversion to insulin might result in the accumulation of proinsulin in *Cnot7*-KO β cells. This prediction was confirmed by immunofluorescence using antibodies against proinsulin and insulin (Figures 4F and 4G). As a consequence of accumulated proinsulin caused by the lack of CNOT7, serum proinsulin was increased in *Cnot7*-KO mice compared to WT (Figure 4H). These data suggested that the conversion of proinsulin to insulin is impaired in *Cnot7*-KO β-cells, leading to the accumulation of proinsulin with reduced mature insulin.

**Figure 4.**
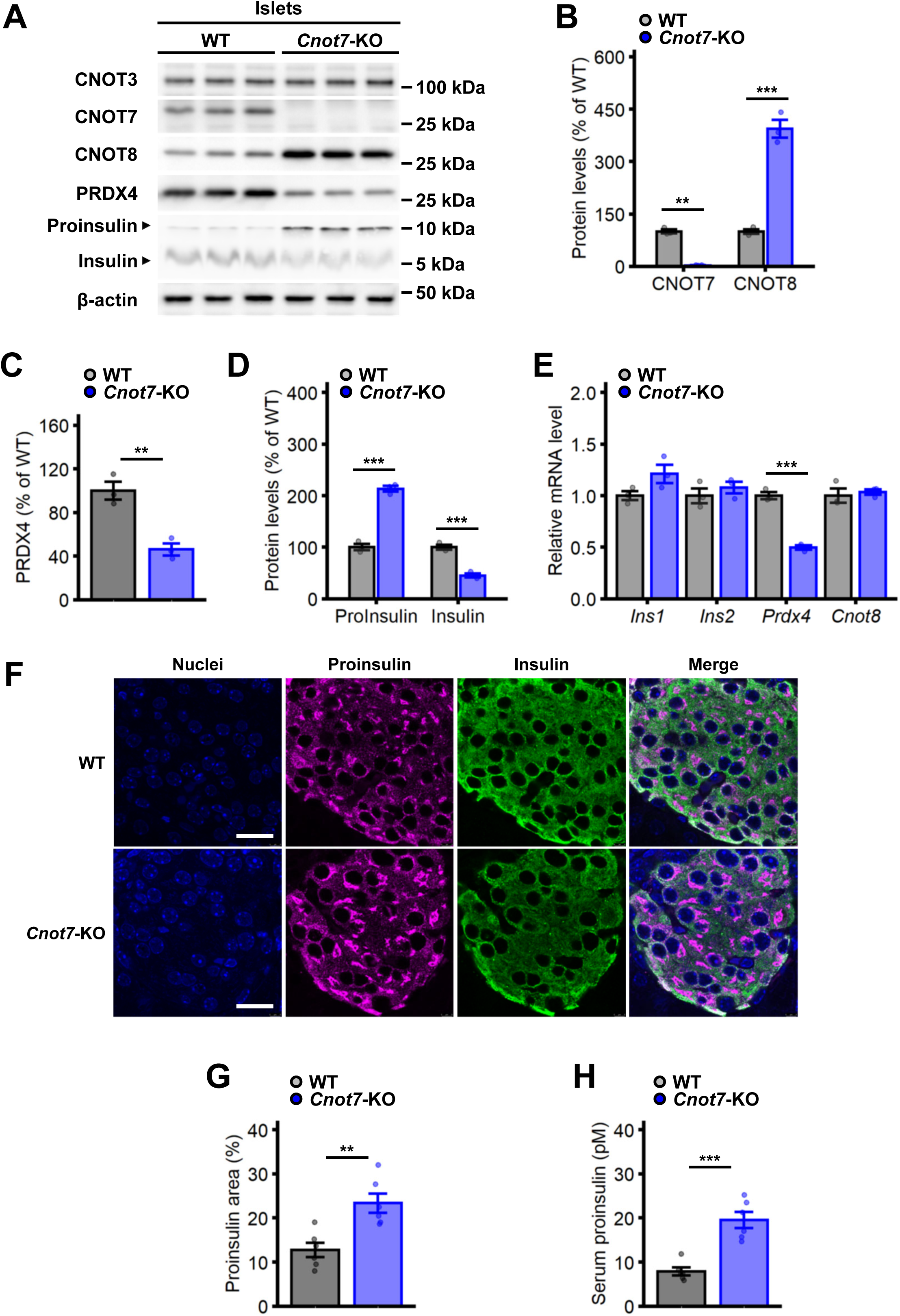
Impaired conversion of proinsulin to insulin in *Cnot7*-KO pancreatic β cells. **(A-D)** Protein expression of CNOT3, CNOT7, CNOT8, PRDX4, proinsulin, insulin and β-actin were determined by western blotting using 8-week-old WT (n=3) and *Cnot7*-KO (n=3) islets. Band intensities of CNOT7 and CNOT8 **(B)**, PRDX4 **(C)** and proinsulin/insulin **(D)** in WT (gray) and *Cnot7*-KO (blue) islets were measured using NIH ImageJ. Average band intensities of WT mice were set as 100%. Data represent means ±SEM. Unpaired Student’s t-test, *p<0.05; **p<0.01; and ***p<0.001. **(E)** Relative mRNA exprssion of *Ins1*, *Ins2*, *Prdx4* and *Cnot8* mRNAs in 8-week-old WT (gray; n=3) and *Cnot7*-KO (blue; n=3) islets. Data represent means ±SEM. One-way ANOVA, Tukey post hoc test, *p<0.05; **p<0.01; and ***p<0.001. **(F)** Proinsulin (magenta) and insulin (green) expression was determined by Immunofluorescence in 8-week-old WT and *Cnot7*-KO pancreatic β cells. Nuclei were stained by DAPI (blue). Merged images are also shown. Scale bar = 20 µm. **(G)** Quantification of proinsulin (magenta) and insulin (green) expression in 8-week-old WT (gray; n=6) and *Cnot7*-KO (blue; n=6) pancreatic β cells. Data represent means ±SEM. One-way ANOVA, Tukey post hoc test, *p<0.05; **p<0.01; and ***p<0.001. **(H)** Serum proinsulin was determined by proinsulin ELISA in 8-week-old WT (gray; n=6) and *Cnot7*-KO (blue; n=6) mice. Data represent means ±SEM. One-way ANOVA, Tukey post hoc test, *p<0.05; **p<0.01; and ***p<0.001.

### MSI2 interacts with *Prdx4* mRNA, contributing to its stability by the CNOT8-containing CCR4-NOT complex

Since CNOT7 is a catalytic subunit of the CCR4-NOT complex, the global length of mRNA poly(A) tails was investigated using 8-week-old WT and *Cnot7*-KO islets by bulk poly(A) assay. There was no significant difference in poly(A) tail length between WT and *Cnot7*-KO islets, implying that CNOT7 and CNOT8 are redundant for most of mRNAs (Figure 5A). Poly(A) tail (PAT) assays showed that the poly(A) tail of *Prdx4* mRNA was shorter in *Cnot7*-KO islets, implying that its poly(A) tail is deadenylated by increased CNOT8 in *Cnot7*-KO islets (Figures 5B and 5C). In contrast, the length of poly(A) tail of *Ins2* mRNA is elongated in *Cnot7*-KO islets by PAT assay, suggesting that CNOT7 contributes to the deadenylation of *Ins2* mRNA (Figures 5D and 5E). To investigate the interaction between mRNAs and the CCR4-NOT complex containing either CNOT7 or CNOT8, which mutually exclusively bind to the same site of the CCR4-NOT complex, RNA immunoprecipitation (RIP) was conducted using anti-CNOT7 or anti-CNOT8 antibodies. In accordance with PAT assay, *Ins2* mRNA interacted with the CNOT7-containing CCR4-NOT complex more than that containing CNOT8, whereas *Prdx4* mRNA interacted with the CNOT8-containing CCR4-NOT complex more than that containing CNOT7 (Figure 5F). Since the CCR4-NOT complex is recruited to its target mRNAs via specific RNA-binding proteins (RBPs), the nucleic acid sequence of *Prdx4* mRNA was examined for potential RBP binding sites (Figure S4). Intriguingly, a conserved Musashi-2 (MSI2)-binding motif was identified in the coding sequence of *Prdx4*, close to the stop codon, implying a direct interaction between MSI2 and *Prdx4* mRNA. To investigate the direct interaction between MSI2 and *Prdx4* mRNA, electrophoretic mobility shift assay (EMSA) was performed using a *Prdx4* RNA oligo containing the MSI2 binding motif and recombinant MSI2 protein. MSI2 interacted with *Prdx4* RNA in a concentration-dependent manner (Figure 5G). This interaction was specific to MSI2 and *Prdx4* RNA, as MSI2 did not interact with a random RNA sequence (*N_20_*) (Figure 5H) and recombinant glutathione S-transferase (GST) did not interact with *Prdx4* RNA (Figure 5I). To confirm that the CCR4-NOT complex containing CNOT8 is recruited to *Prdx4* mRNA via MSI2, *Msi2*-knockdown (KD) MIN6 cells were generated. Protein expression of CNOT3, CNOT7 and CNOT8 was not altered by reduced MSI2 (Figure 5J). *Prdx4* mRNA expression is increased in *Msi2*-KD MIN6 cells (Figure 5K) in accordance with increased stability of *Prdx4* mRNA by reduced MSI2 (Figure 5L), implying that the CCR4-NOT complex is recruited to *Prdx4* mRNA via MSI2 (Figure S4B).

**Figure 5.**
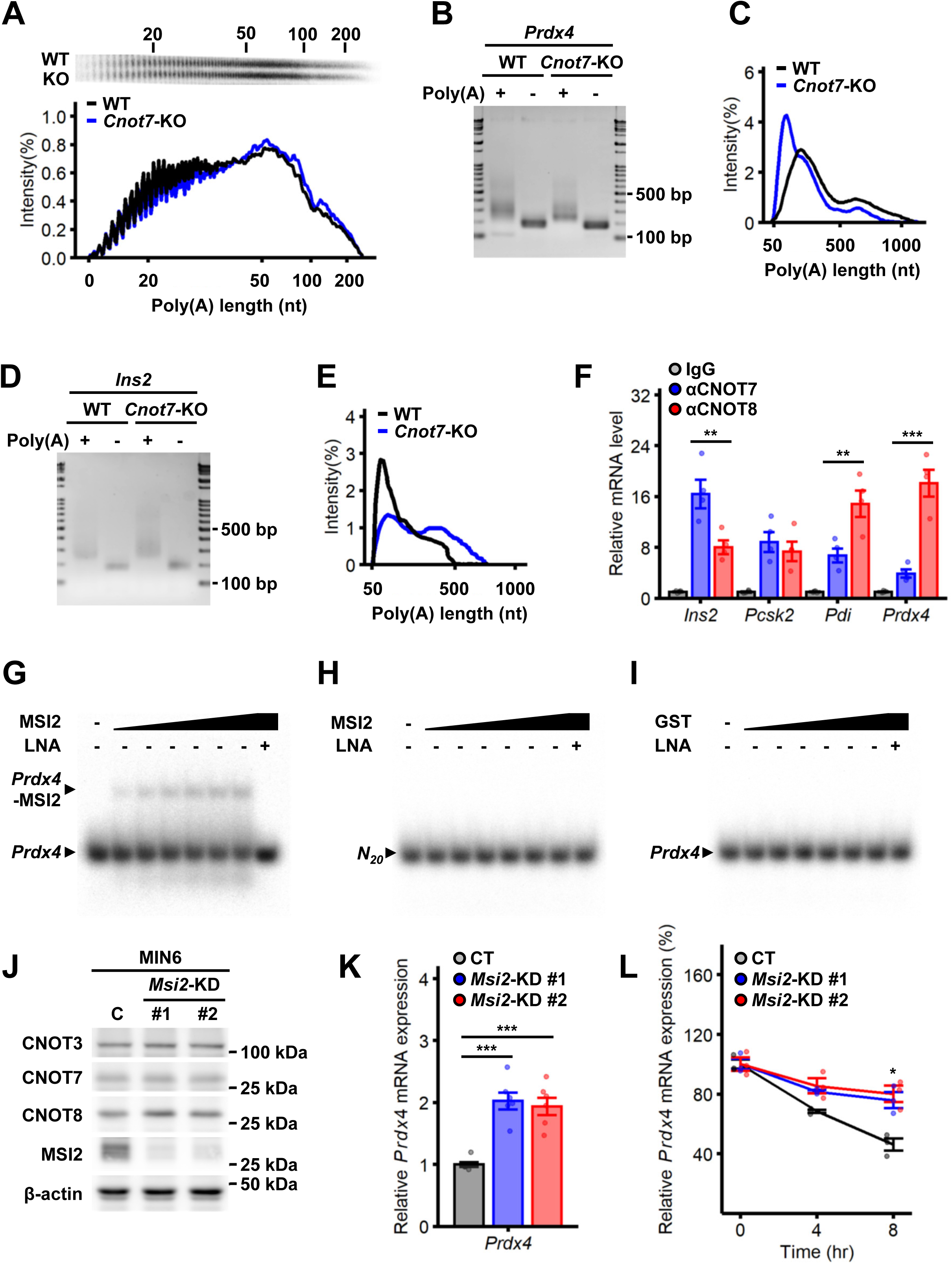
The length of the poly(A) tail of *Prdx4* mRNA is shortened in *Cnot7*-KO islets. **(A)** Poly(A) tail lengths of global mRNAs of 8-week-old WT (black; n=3) and *Cnot7*-KO (blue; n=3) islets were determined by bulk poly(A) assay. Densitograms of poly(A) tail lengths are shown. **(B)** The poly(A) tail length of *Prdx4* mRNA in 8-week-old WT and *Cnot7*-KO islets was determined by PAT assay. PCR products with total RNA treated with RNase H in the presence of oligo(dT) are shown as the length of fragments without poly(A) tails. **(C)** Densitograms of poly(A) tail lengths of *Prdx4* mRNA in WT (black) and *Cnot7*-KO (blue) islets in Figure 5B. **(D)** The poly(A) tail length of *Ins2* mRNA in 8-week-old WT and *Cnot7*-KO islets was determined by PAT assay. PCR products with total RNA treated with RNase H in the presence of oligo(dT) are shown as the length of fragments without poly(A) tails. **(E)** Densitograms of poly(A) tail lengths of *Ins2* mRNA in WT (black) and *Cnot7*-KO (blue) islets in Figure 5D. **(F)** CNOT7-interacting (blue) or CNOT8-interacting (red) mRNAs (*Ins2*, *Pcsk2*, *Pdi* and *Prdx4*) were determined by RNA immunoprecipitation (RIP) using anti-CNOT7 or anti-CNOT8 antibodies, respectively, followed by qPCR. Values of control mouse IgG-interacting mRNAs (gray) are set as 1. Data represent means ±SEM. One-way ANOVA, Tukey post hoc test, *p<0.05; **p<0.01; and ***p<0.001. **(G)** The direct interaction between *Prdx4* mRNA and MSI2 was determined by electrophoresis mobility shift assay (EMSA). Recombinant MSI2 protein was incubated with *Prdx4* RNA oligo containing MSI2-binding motif. Locked nucleic acid (LNA) encoding reverse and complement sequence of MSI2-binding motif was added as a competitor. **(H)** The direct interaction between random sequence RNA oligos (*N_20_*) and MSI2 was determined by EMSA. **(I)** The direct interaction between *Prdx4* RNA oligo and recombinant GST protein was determined by EMSA. **(J)** CNOT3, CNOT7, CNOT8, MSI2 and β-actin were determined by western blotting in control (C) and *Msi2*-knockdown (*Msi2*-KD #1 and #2) MIN6 cells. **(K)** Relative *Prdx4* mRNA expression in control (CT; gray), *Msi2*-KD (KD#1; blue, KD#2; red) MIN6 cells was determined by qPCR. Average value in control cells was set as 1. **(L)** *Prdx4* mRNA stability was determined by qPCR using control MIN6 cells (CT; black; n=3) and *Msi2*-KD (KD#1; blue; n=3, KD#2; red; n=3, respectively) MIN6 cells were treated with actinomycin D for 0, 4 and 8 hours. Average value of control cells at 0 hour was set as 100%. Data represent means ±SEM. Two-way ANOVA followed by Bonferroni post hoc test, *p<0.05; **p<0.01; and ***p<0.001.

### MSI2 interacts with CNOT8 at C-terminus, but not with CNOT7

CNOT7 and its paralog, CNOT8, exhibit high homology between their amino acid sequences, in which a majority of functionally important amino acid residues, such as for catalytic activity (D40) and binding to CNOT6/6L (C67, L71), CNOT1 (M141) and BTG/TOB family proteins (E247, Y260), are conserved. However, distinctive amino acid sequences are present at their C-termini (Figure S5A). To investigate the interaction between MSI2 and the CCR4-NOT complex, FLAG-tagged components of the CCR4-NOT complex (CNOT2, CNOT3, CNOT7 and CNOT8) were co-expressed with HA-tagged MSI2 in HEK293T cells. Co-immunoprecipitation of MSI2 and CNOT7 was greatly reduced compared to the other CCR4-NOT subunits, suggesting that MSI2 preferentially interacts with the CNOT8-containing CCR4-NOT complex (Figures 6A and 6B). Next, the interaction of MSI2 and CNOT7 or CNOT8 in the presence or absence of the CCR4-NOT complex was investigated by immunoprecipitation using anti-FLAG antibody. FLAG-tagged wild-type CNOT7 and CNOT8, and the mutants, in which the CNOT1-binding site was disrupted (M141A), were expressed with HA-tagged MSI2 in HEK293T cells. The interaction between MSI2 and CNOT8-M141A was impaired, revealing that MSI2 preferentially interacts with CCR4-NOT complex-bound CNOT8 more than free CNOT8 (CNOT8-M141A) (Figures 6C and 6D). Since MSI2 selectively interacts with CNOT8 but not the highly similar CNOT7, we hypothesized that MSI2 might interact with CNOT8 at its C-terminus, the region in which CNOT7 and CNOT8 proteins are most different (Figure S5A). To investigate the importance of the C-terminus of CNOT8 for MSI2-binding, FLAG-tagged C-terminal truncated CNOT7 and CNOT8 mutants (CNOT7-D and CNOT8-D) were generated along with the mutants whose C-termini were replaced between CNOT7 and CNOT8 (CNOT7-R and CNOT8-R). These CNOT7/CNOT8 mutants were expressed together with MSI2 in HEK293T cells, and their interaction was analyzed by immunoprecipitation (Figures 6E and 6F). The interaction between MSI2 and CNOT8-D was inhibited, revealing that the C-terminus of CNOT8 is critical for MSI2-binding. Additionally, the interaction between MSI2 and CNOT8-R is less inhibited, suggesting that there might be another crucial MSI2-binding site in CNOT8 other than the C-terminus that might contribute to MSI2-binding together with its C-terminus.

**Figure 6.**
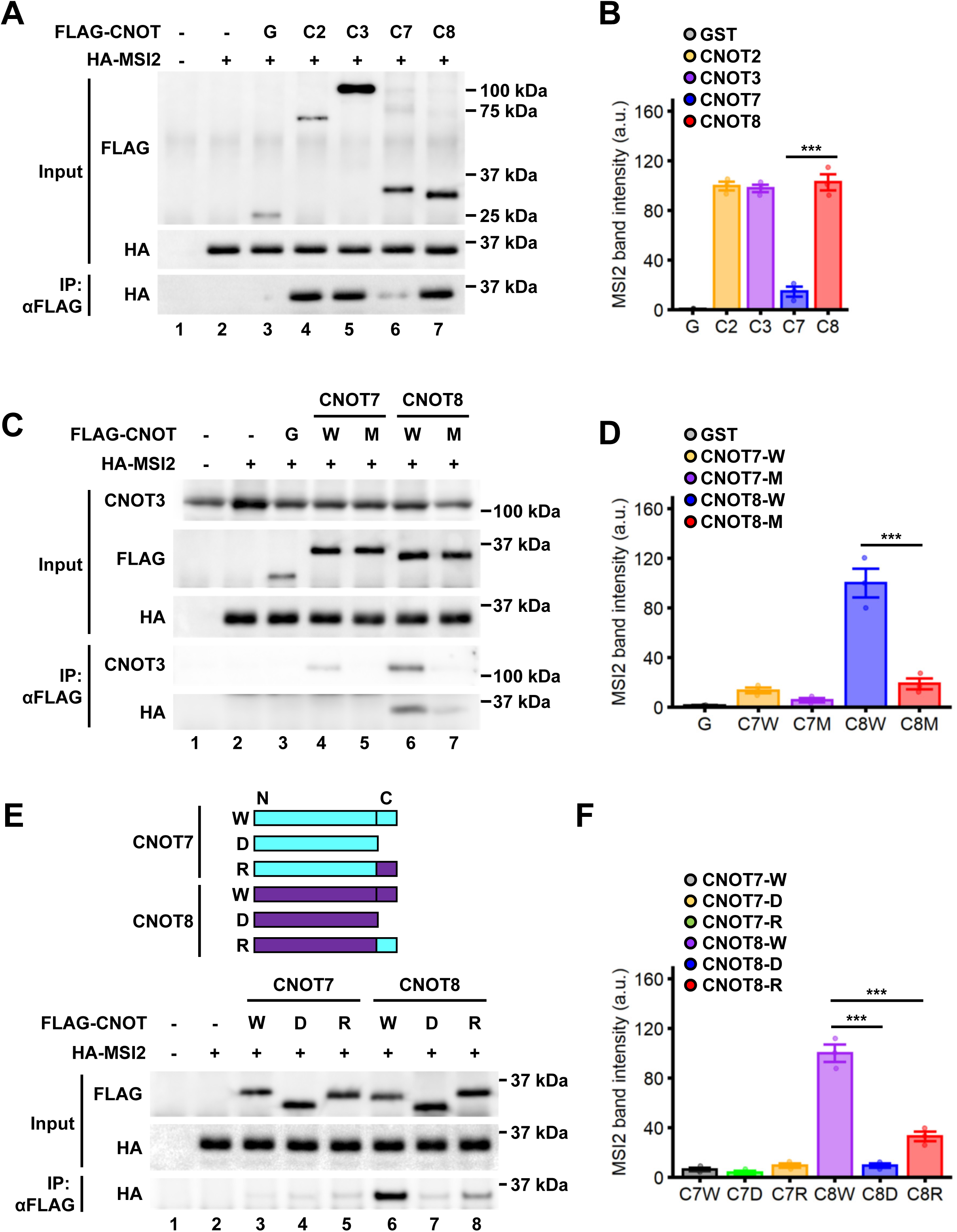
MSI2 interacts with CNOT8 preferentially at its C terminus. **(A)** The interaction between MSI2 and components of the CCR4-NOT complex was determined by immunoprecipitation using anti-FLAG antibody. FLAG-tagged GST (G), CNOT2 (C2), CNOT3 (C3), CNOT7 (C7) and CNOT8 (C8) were expressed along with HA-tagged MSI2 in HEK293T cells as indicated. The top panel represents protein expression of FLAG-tagged proteins, the middle panel represents protein expression of HA-tagged MSI2, and the bottom panel represents immunoprecipitated HA-tagged MSI2 by western blotting. The representative experiment of three independent experiments is presented. **(B)** Quantitative analysis of protein bands in panel 6A. Band intensities (arbitrary units, a.u.) of immunoprecipitated MSI2 against GST (gray; G), CNOT2 (orange; C2), CNOT3 (purple; C3), CNOT7 (blue; C7) and CNOT8 (red; C8) were measured using NIH ImageJ. The average value of band intensities (n=3) of CNOT8-immunoprecipitated MSI2 were set as 100%. Data represent means ±SEM. One-way ANOVA, Tukey post hoc test, *p<0.05; **p<0.01; and ***p<0.001. **(C)** The interaction of MSI2 with either CNOT7 or CNOT8 in the absence or presence of the CCR4-NOT complex was determined by immunoprecipitation using anti-FLAG antibody. FLAG-tagged GST (G), CNOT7-WT (W), CNOT7-M141A (M), CNOT8-WT (W) and CNOT8-M141A (M) were expressed along with HA-tagged MSI2 in HEK293T cells as indicated. From the top to the bottom panels, protein expression of CNOT3, FLAG-tagged proteins, HA-tagged MSI2, immunoprecipitated CNOT3 and HA-tagged MSI2 was determined by western blotting. The representative experiment of three independent experiments is presented. **(D)** Quantitative analysis of protein bands in panel 6C. Band intensities of immunoprecipitated MSI2 against GST (gray; G), CNOT7-WT (orange; C7W), CNOT7-M141A (purple; C7M), CNOT8-WT (blue; C8W) and CNOT8-M141A (red; C8M) were measured using NIH ImageJ. The average value of band intensities (n=3) of CNOT8-WT-immunoprecipitated MSI2 were set as 100%. Data represent means ±SEM. One-way ANOVA, Tukey post hoc test, *p<0.05; **p<0.01; and ***p<0.001. **(E)** The interaction of MSI2 to either CNOT7- or CNOT8-deletion mutants was determined by immunoprecipitation using anti-FLAG antibody. A diagram of CNOT7- (light blue) and CNOT8- (dark purple) deletion mutants is presented on the top. From the top to the bottom, CNOT7-WT (W; a full length), CNOT7-D (D; C-terminal truncated CNOT7), CNOT7-R (R; the C-terminus of CNOT7 was replaced with that of CNOT8), CNOT8-WT, CNOT8-D (D; C-terminal truncated CNOT8) and CNOT8-R (R; the C-terminus of CNOT8 was replaced with that of CNOT7) are shown. FLAG-tagged CNOT7and CNOT8 were expressed along with HA-tagged MSI2 in HEK293T cells as indicated. The top panel represents protein expression of FLAG-tagged proteins, the middle panel represents protein expression of HA-tagged MSI2, and the bottom panel represents immunoprecipitated HA-tagged MSI2 by western blotting. The representative experiment of three independent experiments is presented. **(F)** Quantitative analysis of protein bands in panel 6E. Band intensities of immunoprecipitated MSI2 against CNOT7-WT (gray; C7W), CNOT7-D (green; C7D), CNOT7-R (orange; C7R), CNOT8-WT (purple; C8W), CNOT8-D (blue; C8D) and CNOT8-R (red; C8R) were measured using NIH ImageJ. The average value of band intensities (n=3) of CNOT8-WT-immunoprecipitated MSI2 were set as 100%. Data represent means ±SEM. One-way ANOVA, Tukey post hoc test, *p<0.05; **p<0.01; and ***p<0.001.

In summary, insulin biosynthesis is impaired in *Cnot7*-KO pancreatic β cells by reduced *Prdx4* mRNA, which is destabilized by the CNOT8-containing CCR4-NOT complex via interaction with MSI2 (Figure 7).

**Figure 7.**
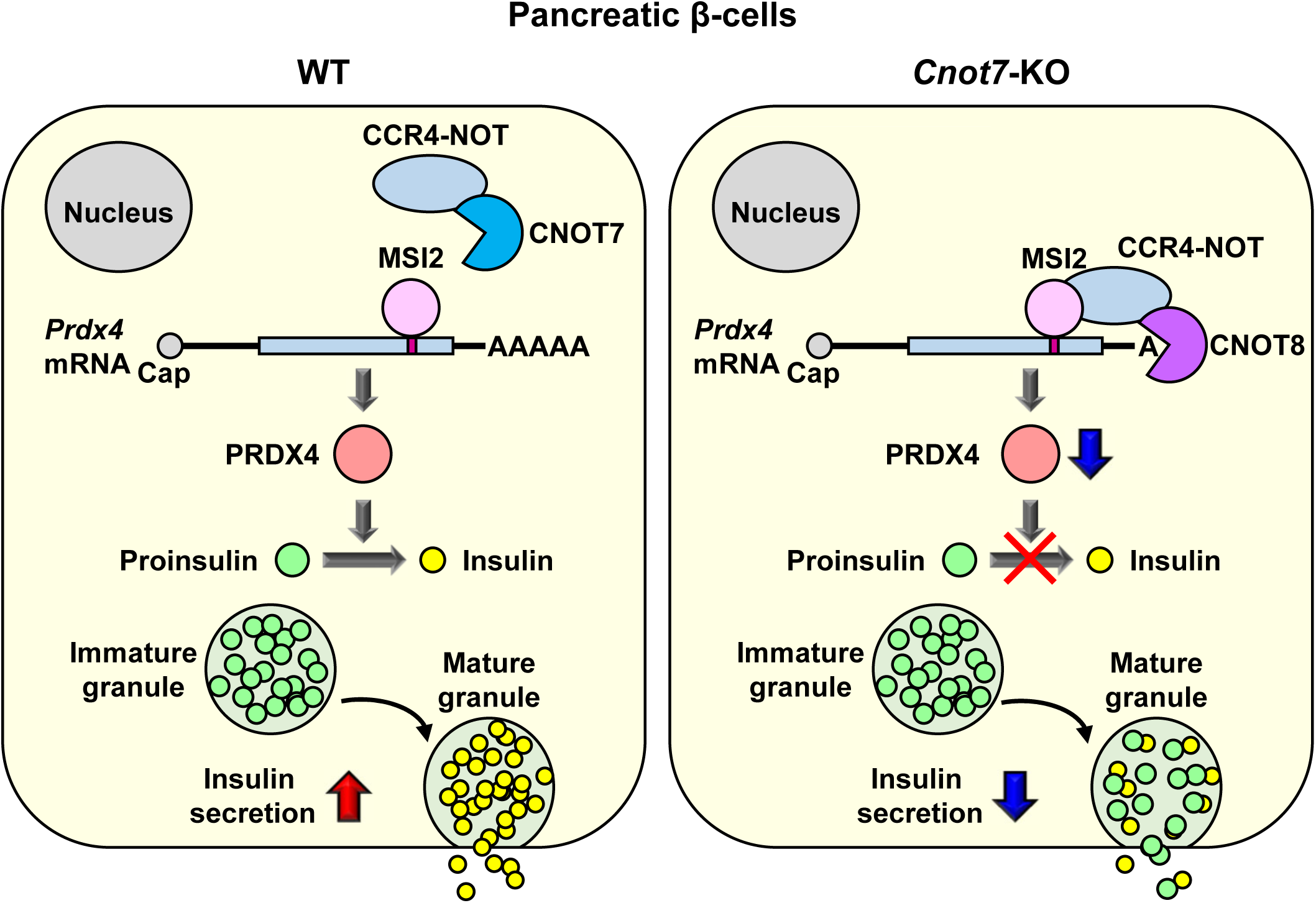
Impaired conversion of proinsulin to insulin by reduced PRDX4 in *Cnot7*-KO pancreatic β cells. PRDX4 is essential for the formation of disulfide bonds inside proinsulin, which is the first step of the conversion to insulin in pancreatic β cells. Insulin biosynthesis in *Cnot7*-KO pancreatic β cells is inhibited by impaired conversion of proinsulin to insulin by reduced PRDX4 caused by *Prdx4* mRNA degradation by elevated CNOT8/MSI2-containg CCR4-NOT complex, leading to reduced insulin content and insulin secretion.

## DISCUSSION

In this study, we describe distinctive functions of CNOT7 and CNOT8 that are essential for insulin biosynthesis by targeting specific mRNAs in pancreatic β cells. Although CNOT7 and CNOT8 are redundant to regulate the stability of a majority of mRNAs (Aslam et al., 2009), it is feasible that CNOT7 and CNOT8 target their specific mRNAs because of their distinctive phenotypes in embryogenesis (Figure S6A) (Mostafa et al., 2020a). Male *Cnot7*-KO mice are infertile with dysregulated sperm formation (Nakamura et al., 2004), in which CNOT8 expression is increased by the lack of CNOT7 in *Cnot7*-KO testis with little change of *Cnot8* mRNA expression (Figures S6B, S6C and C6D). If CNOT7 and CNOT8 were completely redundant, this increased CNOT8 expression could compensate the function of CNOT7 that are essential for spermatogenesis. Increased CNOT8 in *Cnot7*-KO testis fails to maintain proper spermatogenesis, revealing their distinctive physiological functions in spermatogenesis (Figure S6E). Although the amino acid sequences of CNOT7 and CNOT8 are highly similar, they differ at their C-termini, which are intrinsically disordered protein regions (IDPRs) composed of positively and negatively charged amino acids (Figure S5A) (Dunker et al., 2013). IDPRs are characterized by conformational flexibility and structural plasticity and have emerged as an important player to engage in a variety of biological processes (Diella et al., 2008). IDRs participate in diverse cellular functions by interacting with other proteins, in which its activity is often regulated by the post-translational modifications such as phosphorylation (van der Lee et al., 2014) (Iakoucheva et al., 2004) (Collins et al., 2005). Intriguingly, distinctive three-dimensional structures at the C-termini of CNOT7 and CNOT8 are predicted by AlphaFold2, in which any obvious structures are absent in CNOT7 whereas an α-helix is present in CNOT8 (Figure S5B). It is feasible that CNOT7 and CNOT8 might interact with distinctive binding partners via these diverse C-terminal regions, giving them different physiological functions.

Through scrutinization of the *Prdx4* mRNA sequence, we identified MSI2 as an RBP that binds to *Prdx4* mRNA and preferentially interacts with CNOT8, but not CNOT7. MSI2 is known to regulate mRNA stability and translation of target mRNAs in a variety of biological processes such as hematopoietic stem cell (HSC) maintenance, epithelial cell migration and malignant transformation (Park et al., 2014) (Park et al., 2015) (Wang et al., 2015) (Bennett et al., 2016) (Lan et al., 2017). MSI2 is a member of the class A/B heterogeneous nuclear ribonucleoproteins (hnRNP) (Sakakibara et al., 2001) containing two RNA recognition motifs (RRMs) at the N-terminus that mediate binding to target mRNAs. Importantly, MSI2 is expressed in pancreatic β cells and contributes to the regulation of insulin expression, β cell apoptosis and proliferation in response to ER stress caused by diabetes (Szabat et al., 2011). Although the importance of MSI2 in post-transcriptional regulation is well known (Bennett et al., 2016) (Katz et al., 2014) (Rentas et al., 2016) (Sheng et al., 2017) (Duggimpudi et al., 2018), little is known about the role of MSI2 coordinated with the CCR4-NOT complex in mRNA stability. Our study sheds light on the possible post-transcriptional regulation of insulin biosynthesis in pancreatic β cells by MSI2, where it recruits the CNOT8-containing CCR4-NOT complex to *Prdx4* mRNA and promotes its destabilization. Since MSI2 interacts preferentially with the CNOT8-containing CCR4-NOT complex rather than free CNOT8 (Figures 6C and 6D), MSI2 might associate in the vicinity of the binding site between CNOT1 and CNOT8. Further experiments are expected to reveal the coordinated post-transcriptional regulation exerted by MSI2 and the CCR4-NOT complex in a variety of other physiological processes.

PRDX4 is an ER-specific antioxidative peroxidase that oxidases protein disulfide isomerase (PDI) and endoplasmic reticulum oxidase 1 (ERO1), leading to the disulfide bond formation that is essential for oxidative protein folding in ER (Frand and Kaiser, 1998) (Pollard et al., 1998) (Hudson et al., 2015). The oxidation-reduction system plays a pivotal role in insulin biosynthesis in pancreatic β cells by forming disulfide bonds in proinsulin, leading to the proper three-dimensional structure of proinsulin. PRDX4 contributes to pancreatic β functions such as resistance to oxidative stress caused by streptozotocin-induced diabetes (Ding et al., 2010), an enhancement in *Ins1* and *Ins2* mRNA expression and increased insulin secretion and content in insulin producing INS-1E cells (Mehmeti et al., 2014). A recent study revealed the interaction between PRDX4 and proinsulin, leading to normal proinsulin folding (Tran et al., 2020). Despite of the importance of PRDX4 in insulin biosynthesis, little is known about the post-transcriptional regulation of *Prdx4* mRNA. Our study describes the post-transcriptional regulation of the mRNA that is crucial for insulin biosynthesis coordinated by an RNA-binding protein and the CCR4-NOT complex, which might be a potential therapeutic target for diabetes by improving insulin biosynthesis.

## RESOURCE AVAILABILITY

### Lead contact

Further information and requests for resources and reagents should be directed to and will be fulfilled by the Lead Contact, Tadashi Yamamoto.

### Materials availability

Mouse line (*Cnot7^+/-^*) is available from the lead contact upon request.

### Data and code availability

- All data reported in this paper will be shared by the lead contact upon request.
- This paper does not report original code.
- Any additional information required to reanalyze the data reported in this paper is available from the lead contact upon request.

## EXPERIMENTAL MODEL AND SUBJECT DETAILS

### Mice

Mice were maintained under a 12-hr light/12-hr dark cycle in a temperature-controlled (22°C) barrier facility with free access to water and normal diet (NCD, CA1-1, CLEA Japan). *Cnot7*-KO mice were described previously (Nakamura et al., 2004). Heterozygous Cnot7^+/-^ mice were crossed to generate *Cnot7*-KO and WT littermates as controls. Experiments using mice were approved by the Animal Care and Use Committee at the Okinawa Institute of Science and Technology Graduate University (OIST).

### Cell culture

MIN6 cells, HEK293T and HEK293FT cells were cultured in DMEM (11965-092; GIBCO) containing 10% fetal bovine serum (FBS) and 1% penicillin/streptomycin. MIN6 cells were a gift from Dr. Seino a Kobe University Graduate School of Medicine.

### Plasmids

pcDNA3-FLAG-GST, pcDNA3-FLAG-CNOT7, pcDNA3-FLAG-CNOT7-M141A, pcDNA3-FLAG-CNOT7-D (1-263 a.a.), pcDNA3-FLAG-CNOT7-R (CNOT7; 1-263 a.a., CNOT8; 264-292 a.a.), pcDNA3-FLAG-CNOT8, pcDNA3-FLAG-CNOT8-M141A, pcDNA3-FLAG-CNOT8-D (1-263 a.a.), pcDNA3-FLAG-CNOT8-R (CNOT8; 1-263 a.a., CNOT7; 264-285 a.a.) and pcDNA3-HA-MSI2 were generated using pcDNA3 vectors. pCMV-FLAG-CNOT2 and pCMV-FLAG-CNOT3 were generated using pCMV-(DYKDDDDK)-N vector. pGEX-MSI2 vector was generated using pGEX-6P-1 (Cytiva) and cDNA encoding MSI2.

## METHOD DETAILS

### Blood analysis

Fed and fasting blood glucose levels were measured using a glucometer (Glutest Neo alpha Sensor or Glutest Mint Ⅱ, Sanwa Kagaku Kenkyusho); blood was taken from tail vein. After mice were euthanized using isoflurane, blood was collected from the inferior vena cava. Concentrations of serum proinsulin and serum insulin were determined using Rat/Mouse Proinsulin ELISA kit (10-1232-01; Mercodia) and Mouse Insulin ELISA kit (10-1247-01, Mercodia), respectively.

### IPGTT and ITT

To test glucose tolerance, mice were fasted for 16 hours. Mice were injected intraperitoneally with glucose (1 g/kg body weight), and blood glucose levels were measured at 0, 15, 30, 60, 90 and 120 minutes using a glucometer. To test insulin tolerance, fed mice were injected intraperitoneally with insulin (1 U/kg body weight). Blood glucose levels were measured at 0, 15, 30, 60, 90 and 120 minutes using a glucometer.

### Isolation of pancreatic islets

Pancreata were perfused via the bile duct with collagenase solution (1 mg/ml collagenase P [11213865001; Sigma] and 1% Bovine Serum Albumin (BSA) Fraction V [10735086001; Roche] in HBSS [14025076; Thermo Fisher Scientific]). Perfused pancreata were collected and incubated at 37°C for 16 minutes with gentle shaking. Dissociated pancreata were passed through a sieve, and collected by centrifugation at 1,000 x g for 2 minutes. After washing by HBSS supplemented with 0.1 %BSA, islets were separated using a gradient of Histopaque-1077 (10771; Sigma) and RPMI-1640 (11875-093; GIBCO) media supplemented with 10% FBS and 1% penicillin/streptomycin by centrifugation at 1,000 x g for 15 minutes. The boundary layer between the Histopaque and the RPMI containing islets was collected in a culture dish. Islets were collected by hand-picking under a microscope, and cultured in 10% FBS-RPMI for further experiments.

### GSIS

For *in vivo* assays, mice were fasted for 16 hours before intraperitoneal administration of glucose (1 g per kg of body weight). Blood samples were collected 0 and 15 minutes after administration. Serum insulin were determined using a Mouse Insulin ELISA kit (Mercodia). For *ex vivo* assays, isolated islets were cultured in Krebs Ringer buffer (KRB; 140 mM NaCl, 3.6 mM KCl, 0.5 mM NaH_2_PO_4_, 2 mM NaHCO_3_, 1.5 mM CaCl_2_, 0.5 mM MgSO_4_, 10 mM HEPES, 0.25% BSA, pH 7.4) containing 2.8 mM glucose for 1 hour. Culture media were replaced with new KRB buffer containing either 2.8 mM or 16.7 mM glucose for 1 hour. Insulin in culture media was determined using a Mouse Insulin ELISA kit, as for secreted insulin. Islets were lysed with acid lysis buffer (1.5% HCl in ethanol) and sonicated five times. After centrifugation at maximum speed for 5 minutes at 4°C, insulin in supernatant was determined by a Mouse Insulin ELISA kit as for insulin content. Genomic DNA was isolated from the residual islet pellet and islet DNA content was determined. Both secreted insulin and insulin content were normalized to DNA content.

### Immunofluorescence

Pancreata were fixed in 4% paraformaldehyde phosphate buffer solution (163-20145; Wako) overnight at 4°C, and then immersed in 70% ethanol. Fixed pancreata were embedded in paraffin and sectioned at 3 µm. Pancreas sections were rehydrated and boiled in 10 mM sodium citrate buffer (pH 6.0) for antigen retrieval. After blocking in PBS with 0.05% Triton X-100 (PBS-T) containing 5% goat serum for 1 hour at RT, sections were incubated with primary antibodies at 4°C overnight. After washing slides in PBS-T, slides were incubated with secondary antibodies for 2 hours at RT. Slides were then washed in PBS-T and incubated with DAPI. Immunofluorescence was detected by a confocal microscope. Signal intensities were quantified from images using ImageJ software. Antibodies used for immunofluorescence are listed in table S1.7.

### β cell mass

The ratio between β cells and pancreas was determined by measuring insulin area and pancreas area of insulin-stained pancreas sections using NIH Image J. To calculate β cell mass, the ratio between β cell area and pancreas area was then multiplied by the pancreas weight. Five insulin-stained sections were used for β cell mass calculation.

### RNA-seq analysis

Total RNA (100 ng) was used for RNA-seq library preparation with a TruSeq Stranded mRNA Library Prep Kit for NeoPrep (NP-202-1001; Illumina) that allows polyA-oligo(dT)-based purification of mRNA, according to the manufacturer’s protocol with minor modification and optimization as follows. Custom dual index adaptors were ligated at the 5’ and 3’-ends of library, and PCR was performed for 11 cycles. 150 base-pair paired-end read RNA-seq was performed with HiSeq 3000/4000 PE Cluster Kit (PE-410-1001; Illumina) and HiSeq 3000/4000 SBS Kit (300 Cycles) (FC-410-1003; Illumina) on HiSeq4000 (Illumina), according to the manufacturer’s protocol, at a depth of at least 40 million reads. For analysis, RNA-seq data were mapped to the Ensembl GRCm38 *Mus musculus* reference genome using tophat2 software (v2.1.1) on Deigo computing cluster at OIST. Mapped reads were sorted with samtools (v1.3.1), then read counts were quantified with HTSeq-count (v0.9.1). Counts were analyzed for differential gene expression using the edgeR package (v3.28.1) by Bioconductor (v3.14) on R (v3.6.3). Differentially expressed genes with the P value < 0.01 were considered as significantly different using three biological replicates. Among significantly different genes, genes with fold change (FC) < 0.67 were considered as down-regulated genes whereas those with FC > 1.5 were considered as up-regulated genes. GO term enrichment analysis was performed using DAVID (david.ncifcrf.gov). GO terms with false discovery rate (FDR) < 0.05 were considered significantly enriched.

### Mass spectrometry

Protein samples were reduced with dithiothreitol and then alkylated with iodoacetamide. Overnight digestion with Lys-C/Trypsin combo (1:50 enzyme:protein; Promega). After stopping the digestion with 1% trifluoroacetic acid, the peptide mixture was cleaned with desalting C18 tips (StageTip, ThermoFisher Scientific), and subsequently vacuum-centrifuged to dryness, and re-suspended for LC/MS analysis. For data acquisition, data were collected using an Orbitrap Lumos mass spectrometer (ThermoFisher Scientific) coupled with Waters nanoACQUITY liquid chromatography system (Waters Co.). A trap-column (nanoACQUITY UPLC 2G-V/M Trap 5µm Symmetry C18, 180 µm x 20 mm, Waters) and analytical column (nanoACQUITY UPLC HSS T3 1.8 um, 75 µm x 150 mm, Waters Co.) were used for sample chromatographic separation. Peptides were separated at a flow rate of 500 nl/min using a gradient of 1–32% ACN (0.1% formic acid) over 60 min. The CHOPIN method was used on Orbitrap Lumos mass spectrometer using Xcalibur (v.3.0; Thermo Fisher Scientific) without modifications (Davis et al., 2017). For data analysis, raw data files were searched against a composite target/decoy database using SEQUEST from Proteome Discoverer software (PD, v.2.2, ThermoFisher Scientific). The proteome reference UP000000589 database, Mus musculus (Mouse) (Strain: C57BL/6J), was downloaded from UniProt, and combined with contaminants database (cRAP, thegpm.org/crap). MS2 spectra were searched with ± 20 ppm for precursor ion mass tolerance, ± 0.1 Da for fragment ion mass tolerance, using Trypsin as enzyme, two missed cleavages maximum, dynamic modifications for oxidation of methionine and deamidation of glutamine and asparagine, fixed modification carbamidomethylation of cysteine residues. Label-free quantification based on peak intensity, using unique and razor peptides, and normalized using specific protein amount.

### Immunoblotting

Cells were washed with ice-cold PBS and lysed in lysis buffer (50 mM Tris-HCl [pH 7.5], 100 mM KCl, 1 % Triton X-100, 20 mM β-glycerolphosphate, 1 mM DTT, 0.1 mM Na_3_VO_4_, 5 mM NaF, 1 mM EDTA and 1 mM PMSF) supplemented with EDTA-free Protease Inhibitor Cocktail (Nacalai Tesque). Cells were lysed by three freeze-thaw cycles, and centrifugated at maximum speed for 5 minutes to remove cell debris. Samples were subjected to SDS polyacrylamide gel (SDS-PAGE) electrophoresis, and blotted on 0.2 μm nitrocellulose membranes (GE10600001; Merck). After blocking, membranes were probed with primary antibodies and subsequently incubated with horseradish peroxidase (HRP)-conjugated secondary antibodies. Chemiluminescent signals were detected using an ImageQuant LAS 4000 (GE Healthcare). The detail information about primary and secondary antibodies are indicated in table S1.7.

### Quantitative PCR

cDNA was synthesized using total RNA (1μg), oligo(dT)_12-18_ primer (Invitrogen) and SuperScript Reverse Transcriptase III (Thermo Fisher Scientific) for qPCR using TB Green Premix Ex Taq (Takara) on a ViiA 7 Real-Time PCR system (Applied Biosystems). Primer sequences to detect mRNAs and pre-mRNAs are shown in Tables S1.2 and S1.3, respectively. Target mRNA expression was normalized to *Actb* mRNA.

### Bulk poly(A) assay

Total RNA was labeled with [^32^P]pCp (cytidine 39,59-bis[phosphate]; NEG019A; PerkinElmer) using T4 RNA ligase 1 (M0204S; New England Biolabs) at 16°C overnight. Labeled RNAs were incubated at 85°C for 5 min and placed on ice. Labeled RNAs were digested with RNase A (EN0531; Thermo Fisher Scientific) and RNase T1 (1000 U/µL) (EN0542; Thermo Fisher Scientific) at 37°C for 2 hours in digestion buffer (10 mM Tris–HCl [pH 7.5] and 0.3M NaCl). After adding stop solution (2 mg/ml Proteinase K, 0.025 mM EDTA, and 0.5% SDS), reactions were subsequently incubated at 37°C for 30 minutes. After adding RNA precipitation buffer (0.5 M NH_4_OAc and 10 mM EDTA), digested RNA samples were purified by phenol-chloroform extraction and isopropanol precipitation. RNA was subjected to 8M urea–10% polyacrylamide denaturing gel and autoradiography using Typhoon FLA 9500 Fluorescence Imager (GE Healthcare). Band intensities were quantified using NIH Image J.

### PAT assay

The poly(A) tail length was determined by Poly(A) Tail-Length Assay Kit (764551KT; Affymetrix) according to the manufacturer’s instructions. The first round of PCR was performed using the first gene-specific forward primer and the universal reverse primer. The resulting PCR product was diluted 1:100, and diluted PCR product was used as a template for the second round of PCR using the second gene-specific forward primer and the universal reverse primer. PCR products were subjected to 2% agarose TAE gel. Primers used for PAT assay are listed in Table S1.4.

### Recombinant protein

Rosetta (DE3) competent cells (Novagen) were transformed with pGEX-MSI2, and cultured at 25°C in a shaker. After adding isopropyl β-D-1-thiogalactopyranoside (IPTG) for 2 hours, cells were collected by centrifugation and lysed in lysis buffer (0.1% Triton X-100, 100 µg/ml Lysozyme, 0.05% β-mercaptoethanol in PBS [pH 7.4]) supplemented with protease inhibitors. Cell lysate was sonicated and clarified by a centrifugation. The resultant supernatant was incubated with Glutathione Sepharose 4B (17-0756-01, Cytiva) overnight at 4°C. After washing the beads with PBS containing 0.1% Triton X-100, recombinant MSI2 protein was eluted by elution buffer (50 mM Tris [pH7.0], 150 mM NaCl, 1 mM EDTA, 1 mM DTT, 0.01% Triton-X-100) supplemented with PreScission protease (27-0843-01; Cytiva). MSI2 protein was dialyzed in a dialysis buffer (20 mM HEPES [pH8.0], 50 mM NaCl, 200 µM EDTA, 1 mM DTT, 10% glycerol).

### EMSA

*Prdx4* RNA oligo was labeled with [^32^P]ATP (adenosine 5’-triphospate, [γ-^32^P]; NEG002A; PerkinElmer) using T4 polynucleotide kinase (M0201; New England Biolabs) at 37°C for 30 minutes. [^32^P]ATP-labeled *Prdx4* RNA oligo (0.2 nM) and recombinant MSI2 were incubated at 30°C for 30 minutes in binding buffer (20 mM HEPES-KOH [pH 7.5], 100 mM KCl,10% glycerol, 3 mM MgCl_2_, 1 mM DTT, 0.1% Triton X-100 and 0.1 mg/ml acetylated BSA). The reaction mixtures were supplemented with sample buffer and analyzed on a nondenaturing polyacrylamide gel (7% acrylamide, 0.187% bis/acrylamide, 5% glycerol in 0.5x TBE buffer). The gel was pre-run at 100 V at 4°C for 1 hour prior to sample loading. Electrophoretic separation was performed at 100 V at 4°C. Autoradiography was performed using Typhoon FLA 9500 Fluorescence Imager (GE Healthcare), and band intensities were quantified using NIH Image J.

### Gene silencing using shRNA

Lentivirus expressing non-target shRNA (SHC002) and shRNAs targeting *Msi2* were generated according to manufacturer’s instruction (Sigma). HEK293FT cells were co transfected with vectors expressing shRNA and lentivirus packaging plasmids. MIN6 cells were infected with lentivirus expressing shRNA to generate control and *Msi2*-KD MIN6 cells. shRNAs used for silencing are indicated in table S1.5.

### RNA immunoprecipitation

Cells were lysed in 1 ml TNE buffer (50 mM Tris-HCl [pH 7.5], 150 mM NaCl, 1 mM EDTA, 1% NP40 and 1 mM PMSF) supplemented with protease inhibitor cocktail (03969-21; nacalai tesque) and RNaseOUT Ribonuclease Inhibitor (10777019; ThermoFisher Scientific). After centrifugation at 20,000 x g for 10 minutes, cell lysates were divided to two aliquots at the ratio of 1:10, in which total RNA was extracted from smaller portion by TRIzol Reagent (15596018; ThermoFisher) to determine input mRNAs. The larger portion of cell lysates was incubated with antibodies against proteins of interest or normal mouse IgG at 4°C with rotating for 2 hours. Lysates were then added with Protein G-conjugated Dynabeads (10004D, Invitrogen) at 4°C for 1 hour. After washing beads four times using TNE buffer, beads-associated RNA was extracted using TRIzol Reagent. Relative mRNA expression of input and immunoprecipitated RNA was determined by qPCR.

### Immunoprecipitation

Cell lysates were incubated with anti-FLAG antibody along with Protein G-conjugated Dynabeads (10004D, Invitrogen) at 4°C for 4 hours. Immunoprecipitated proteins were washed four times with lysis buffer, dissolved in Laemmli sample buffer, and analyzed by western blotting.

## QUANTIFICATION AND STATISTICAL ANALYSIS

Statistical analyses were conducted using an unpaired two-tailed Student’s t test, one-way ANOVA followed by Tukey *post hoc* test or two-way ANOVA followed by Bonferroni *post hoc* test as indicated in the figure legends.

## ACKNOWLEDGMENTS

This work was supported by the Okinawa Institute of Science and Technology (OIST) subsidiary budget to TY. AY was supported by a Grant-in-Aid for Scientific Research (C) (18K06975) and a Grant-in-Aid for Scientific Research in Innovative Areas (20H05351), from the Japan Ministry of Education, Culture, Sports, Science and Technology. PS was supported by a Grant-in-Aid for Young Scientist (18K16218) We thank OIST for generous support to the Cell Signal Unit.

## AUTHOR CONTRIBUTIONS

AY conceived the project, designed the experiments, conducted experiments, analyzed the data and wrote the manuscript. PS, ST, YT and RH supported experiments. AVB performed MS analysis. AY performed RNA-seq analysis using a pipeline by Linux and R scripts. GV generated a pipeline for RNA-seq analysis. AY, ST and NF generated graphs by R studio. AY, ST and NF performed statistical analysis using R scripts. AY, PS and TY revised the manuscript.

## SUPPLEMENTAL INFORMATION

**Table S1.1.**
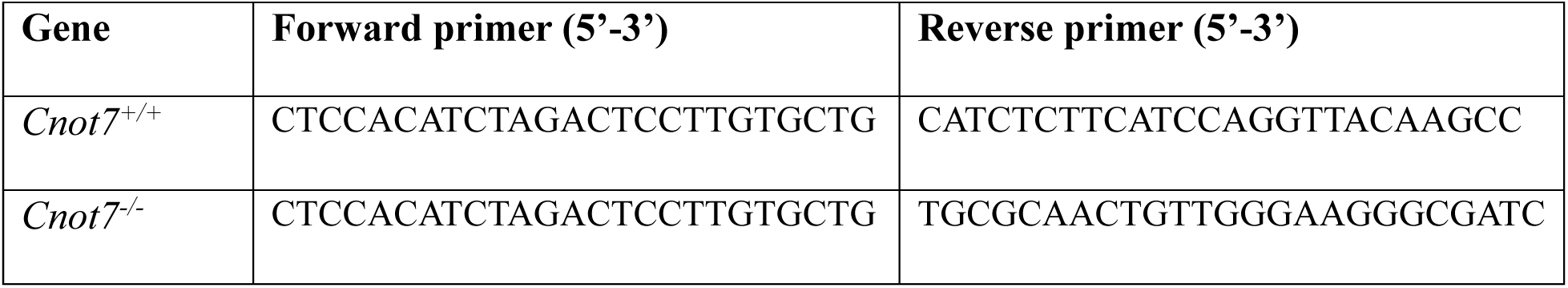
Primers for genotyping of *Cnot7*-KO mice, related to STAR Methods.

**Table S1.2.**
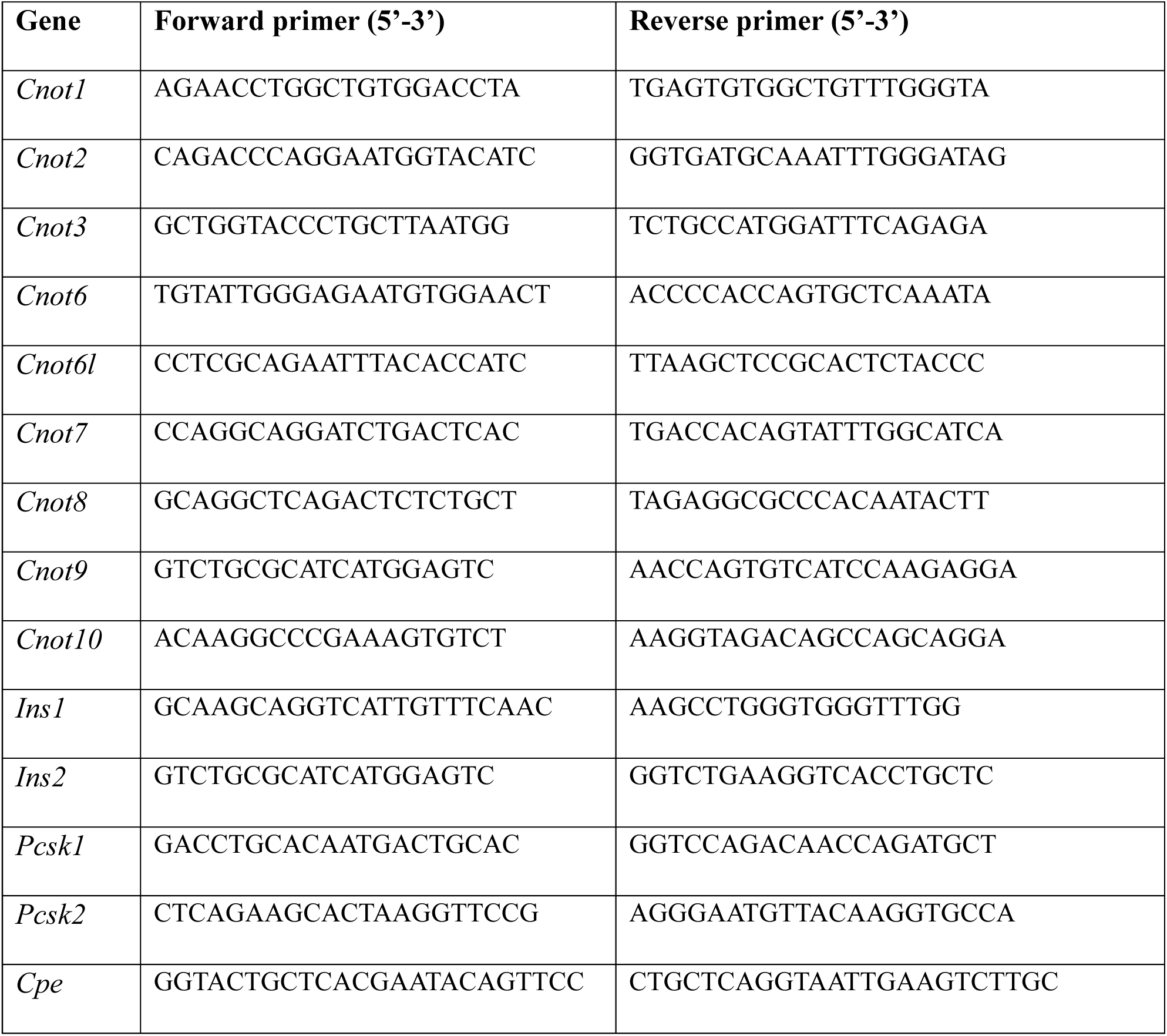
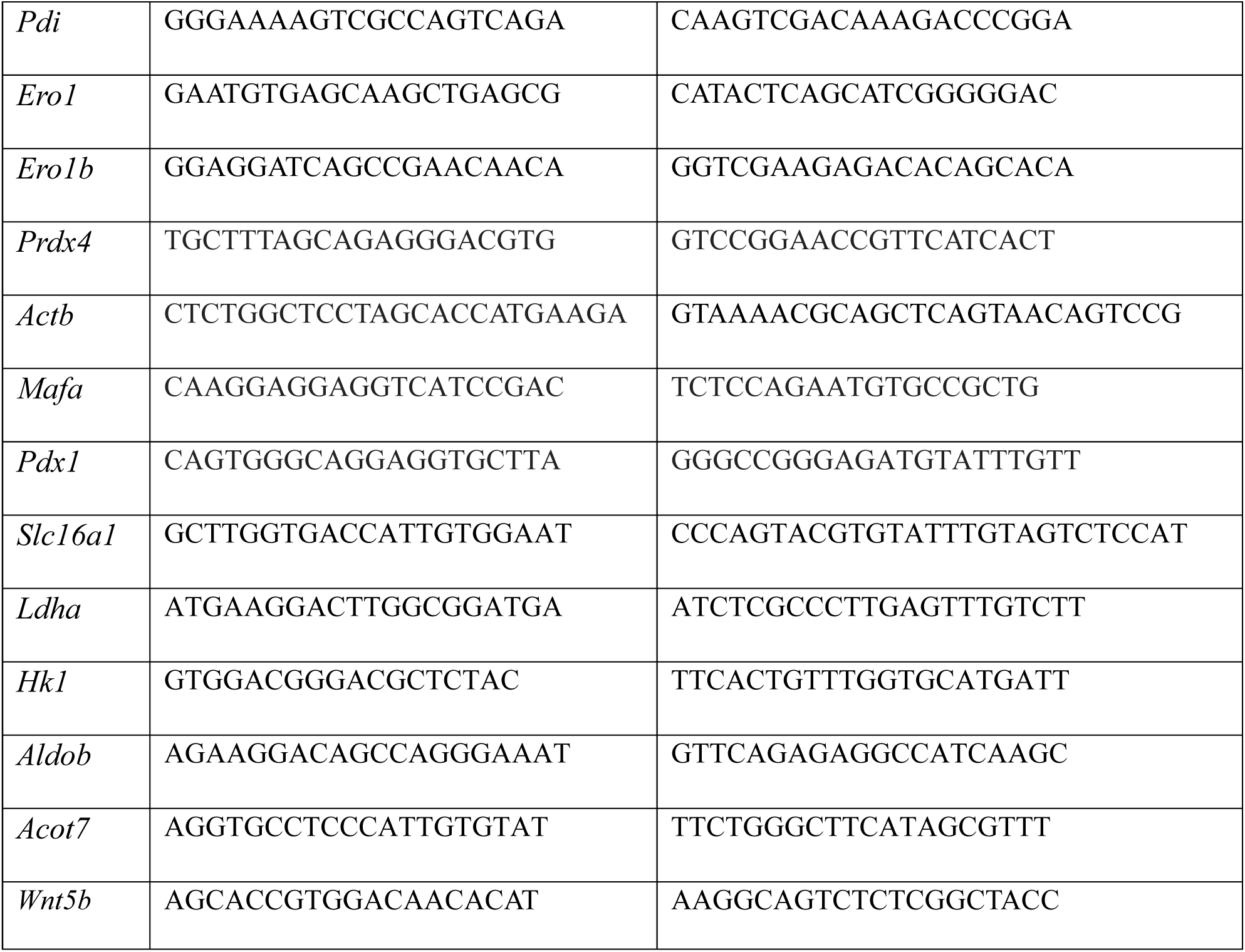
Primers used for qPCR to detect mRNAs, related to STAR Methods.

**Table S1.3.**
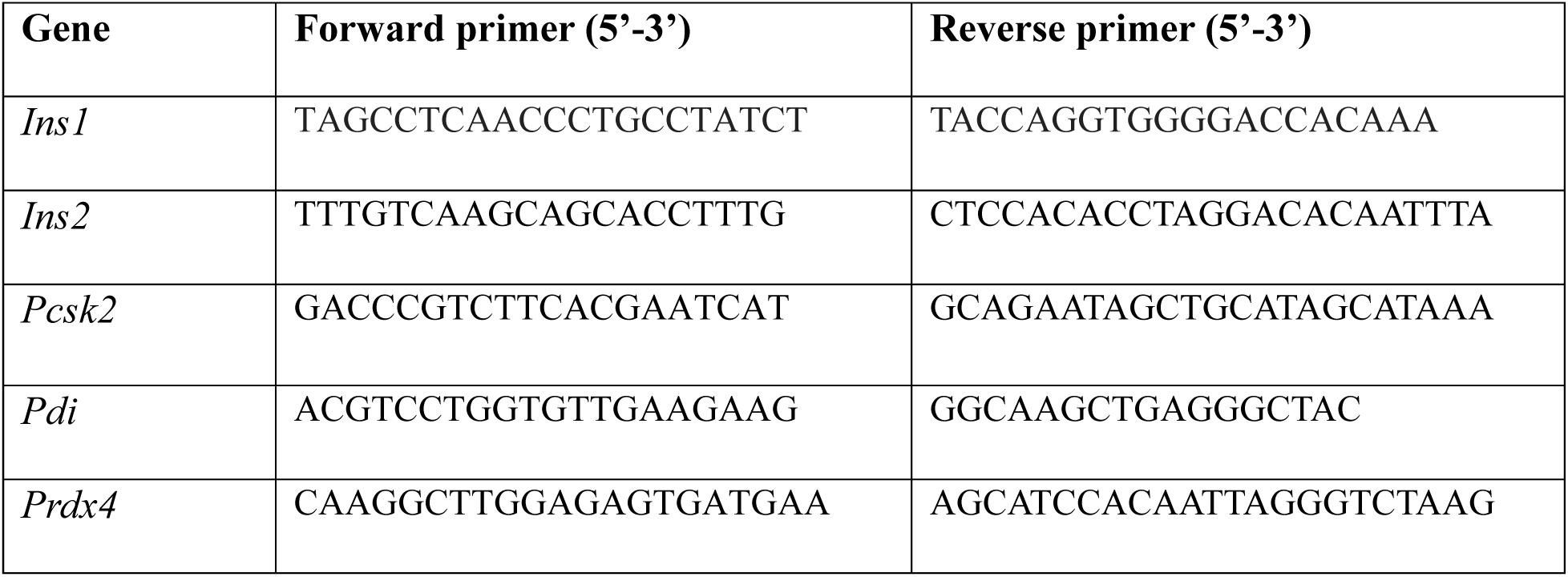
Primers used for qPCR to detect pre-mRNAs, related to STAR Methods.

**Table S1.4.**
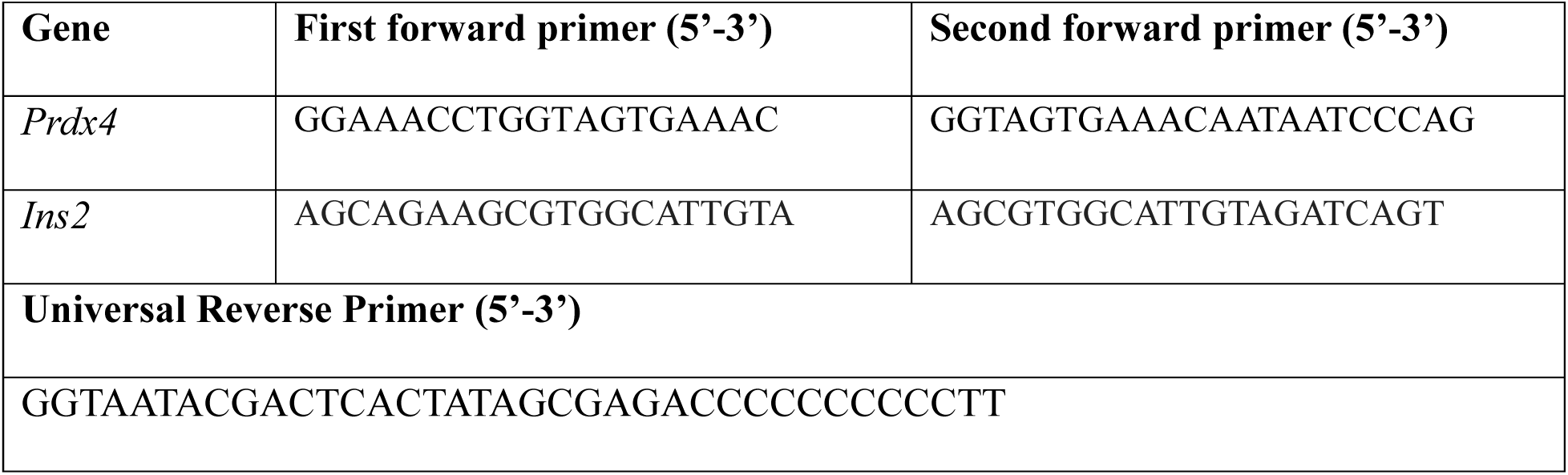
Primers used for PAT assay, related to STAR Methods.

**Table S1.5.**
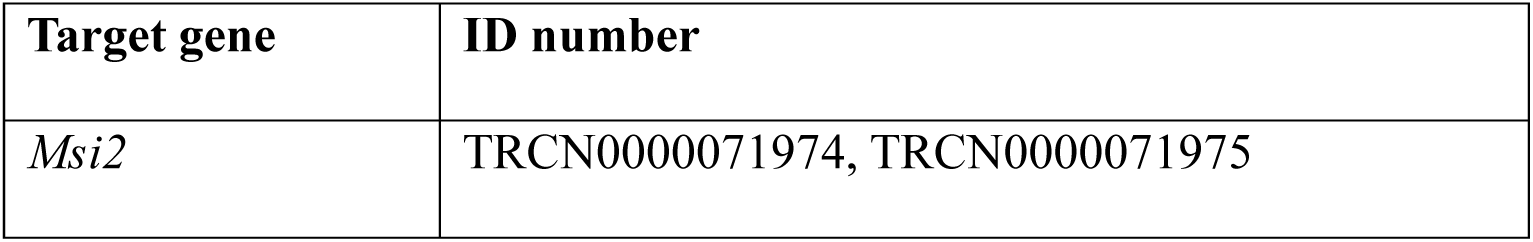
shRNAs used for gene silencing, related to STAR Methods.

**Table S1.6.**
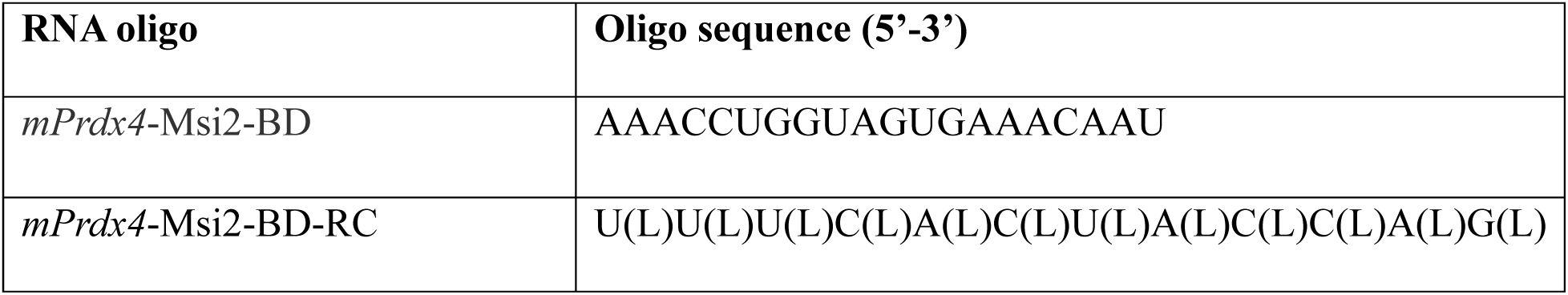
RNA oligos used for EMSA, related to STAR Methods.

**Table S1.7.** Antibodies, related to STAR Methods.

| Antibody | Cat# | Source |
| --- | --- | --- |
| Anti-PRDX4 antibody | 10703-1-AP | Proteintech |
| Anti-MSI2 antibody | Ab76148 | Abcam |
| Anti-FLAG antibody | F1804 | SIGMA |
| Anti-FLAG antibody | PM020 | MBL |
| Anti-HA antibody | # 561 | MBL |
| Anti-CNOT3 antibody | #54 | Biomatrix |
| Anti-CNOT7 antibody | #30B | Biomatrix |
| Anti-CNOT7 antibody | H00029883-M01A | Abnova |
| Anti-CNOT8 antibody | #419 | Biomatrix |
| Anti-CNOT8 antibody |  | This paper |
| Anti-Insulin antibody | #3014 | Cell signaling |
| Anti-Insulin antibody | IR00261 | Agilent |
| Anti-Proinsulin antibody | MAB13361 | R&B systems |
| Anti-Glucagon antibody | Ab92517 | Abcam |
| Anti-β-Catenin antibody | #610153 | BD Laboratories |
| Anti-β-Actin antibody | #4970 | Cell signaling |
| Normal mouse IgG | sc-2025 | Santa Cruz |

---

**Figure S1.**
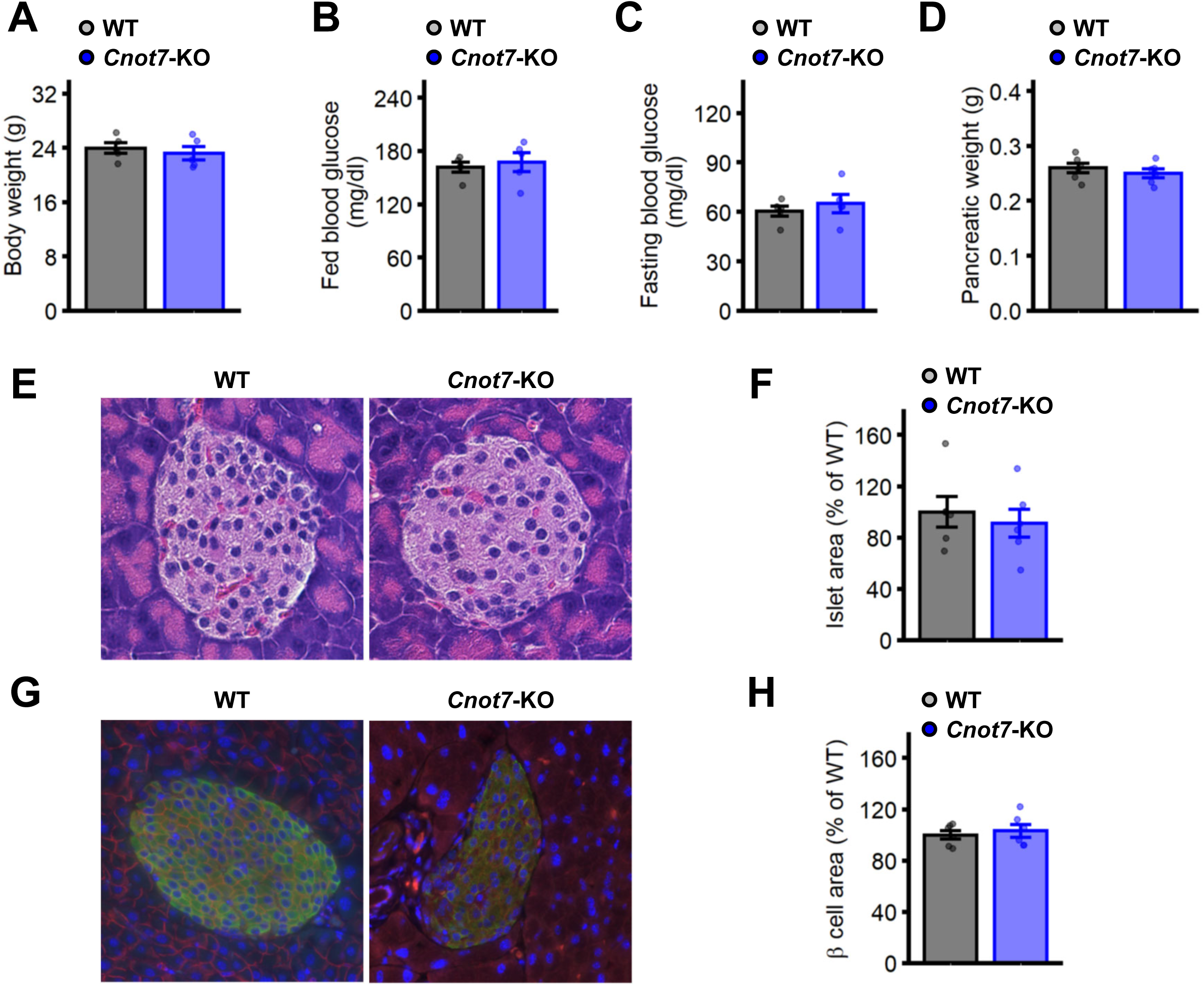
Metabolic phenotypes of *Cnot7*-KO mice, related to Figure 1. **(A-D)** Body weights **(A)**, fed blood glucose **(B)**, fasting blood glucose **(C)** and pancreatic weights **(D)** of 8-week-old *Cnot7*-KO mice (blue; n=6) as compared to WT mice (grey; n=6). Values of the average of WT mice were set as 100 %. Data represent means ±SEM. One-Way ANOVA, post-hoc Tukey test, *p<0.05; **p<0.01; and ***p<0.001. **(E)** Morphology of pancreatic islets of 8-week-old WT and *Cnot7*-KO mice, analyzed by HE staining. **(F)** Islet area of pancreatic sections by HE staining in Figure S1E of 8-week-old WT (grey; n=6) and *Cnot7*-KO (blue; n=6). The average of islet area of WT mice was set as 100%. Data represent means ±SEM. **(G)** Immunohistochemistry of nuclei (blue), insulin (green) and β-catenin (red) of 8-week-old WT and *Cnot7*-KO pancreata. **(H)** Pancreatic β cell area of pancreatic sections by immunohistochemistry in Figure S1G of 8-week-old WT (grey; n=6) and *Cnot7*-KO (blue; n=6). The average of β cell area of WT mice was set as 100%. Data represent means ±SEM.

**Figure S2.**
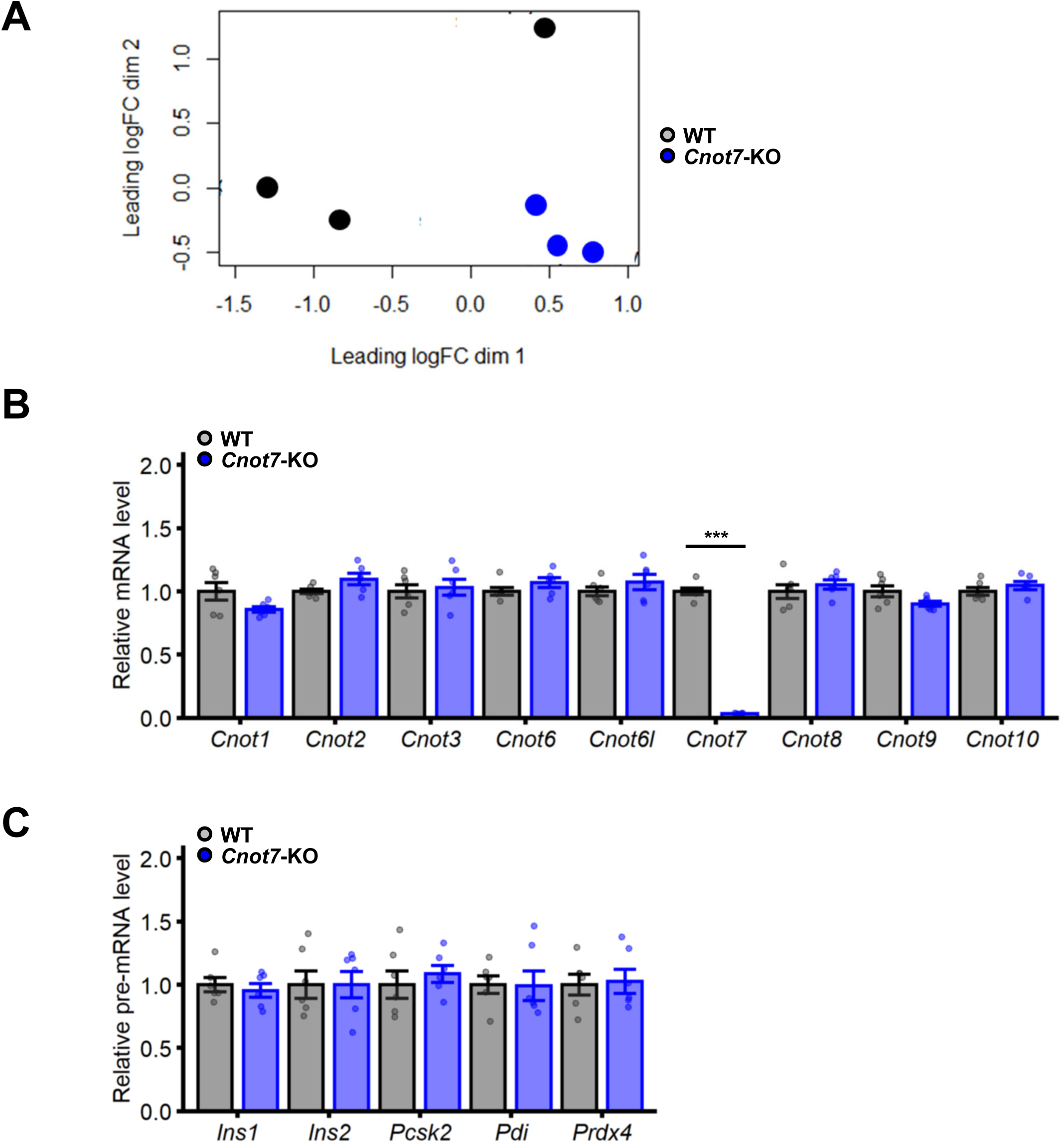
Relative mRNA expression in *Cnot7*-KO islets, related to Figure 2. **(A)** A MDS plot of RNA-seq analysis to show the variation between 8-week-old WT (black; n=3) and *Cnot7*-KO (blue; n=3) islets. **(B)** Relative mRNA expression of *Cnot1*, *Cnot2*, *Cnot3*, *Cnot6*, *Cnot6l*, *Cnot7*, *Cnot8*, *Cnot9* and *Cnot10* in WT (grey; n=6) and *Cnot7*-KO islets (blue; n=6) was determined by qPCR. The values of WT mice were set as 1. Data represent means ±SEM. One-Way ANOVA, post-hoc Turkey test, *p<0.05; **p<0.01; and ***p<0.001. **(C)** Relative pre-mRNA expression of *Ins1*, *Ins2*, *Pcsk2*, *Pdi* and *Prdx4* in WT (grey; n=6) and *Cnot7*-KO islets (blue; n=6) was determined by qPCR. The values of WT mice were set as 1. Data represent means ±SEM.

**Figure S3.**
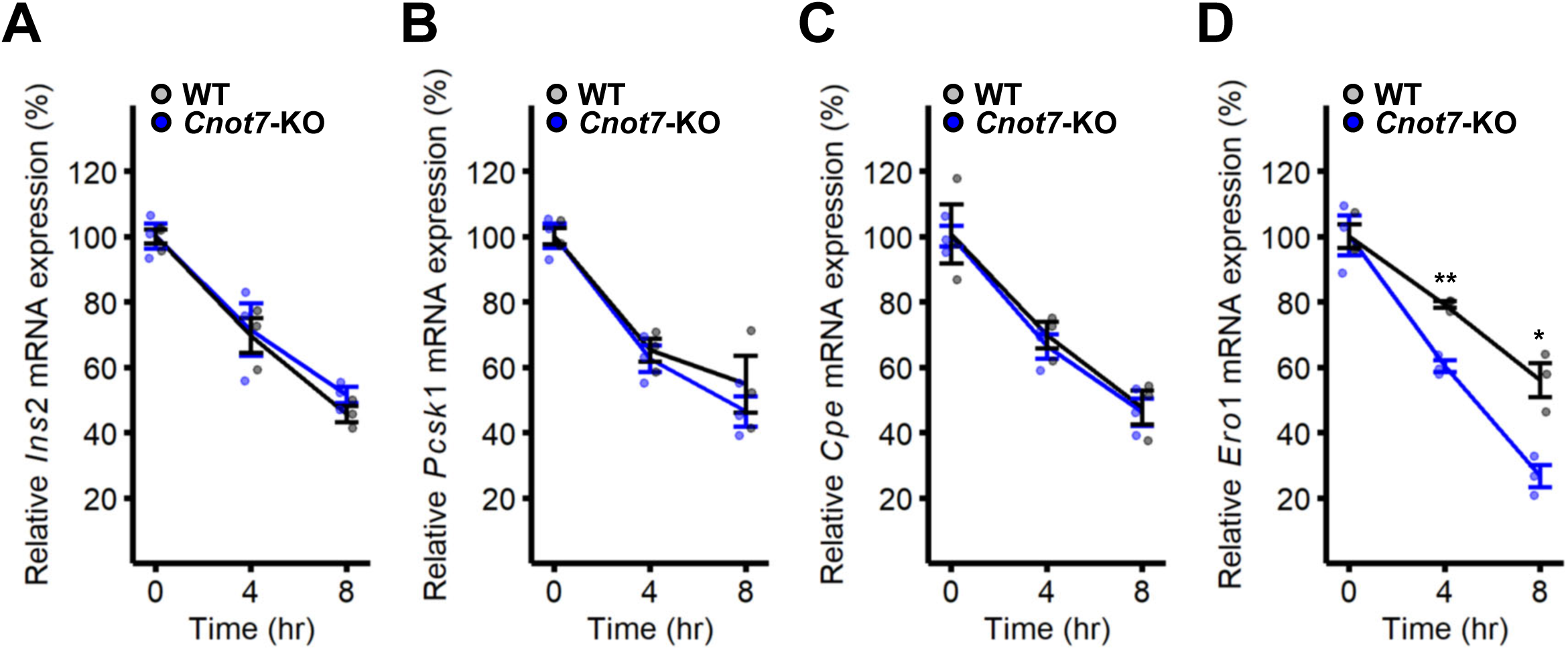
mRNA stabilities in *Cnot7*-KO pancreatic islets, related to Figure 3. **(A-D)** Stabilities of *Ins2* **(A)**, *Pcsk1* **(B)**, *Cpe* **(C)** and *Ero1* **(D)** mRNAs in WT (black, n=3) and *Cnot7*-KO (blue, n=3) pancreatic islets, analyzed by qPCR. Isolated islets were treated by actinomycin D for indicated times. The average values at 0 hour were set as 100%. Data represent means ±SEM. Two-way ANOVA followed by Bonferroni post hoc test, *p<0.05; **p<0.01; and ***p<0.001.

**Figure S4.**
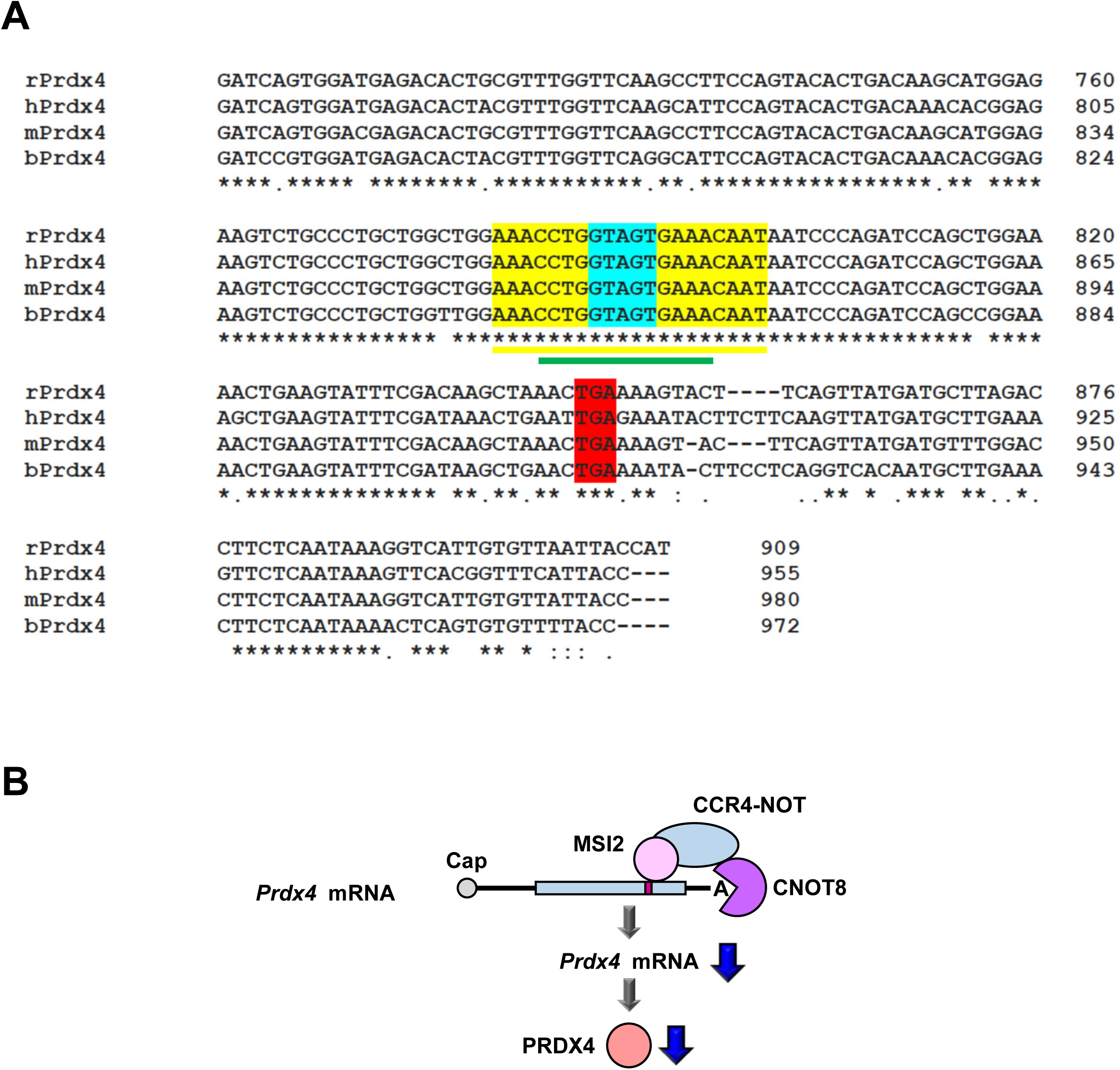
Multiple sequence alignment of *Prdx4* between species, related to Figure 5. **(A)** Nucleic acid alignment between rat (r), human (h), mouse (m) and bovine (b) *Prdx4* mRNAs. MSI2-binding motif (light blue), stop codon (red) and RNA oligos used for EMSA (sense: yellow and anti-sense: green) are shown. **(B)** A diagram of the post-transcriptional regulation of PRDX4 by the interaction between *Prdx4* mRNA, MSI2 and CCR4-NOT complex containing CNOT8.

**Figure S5.**
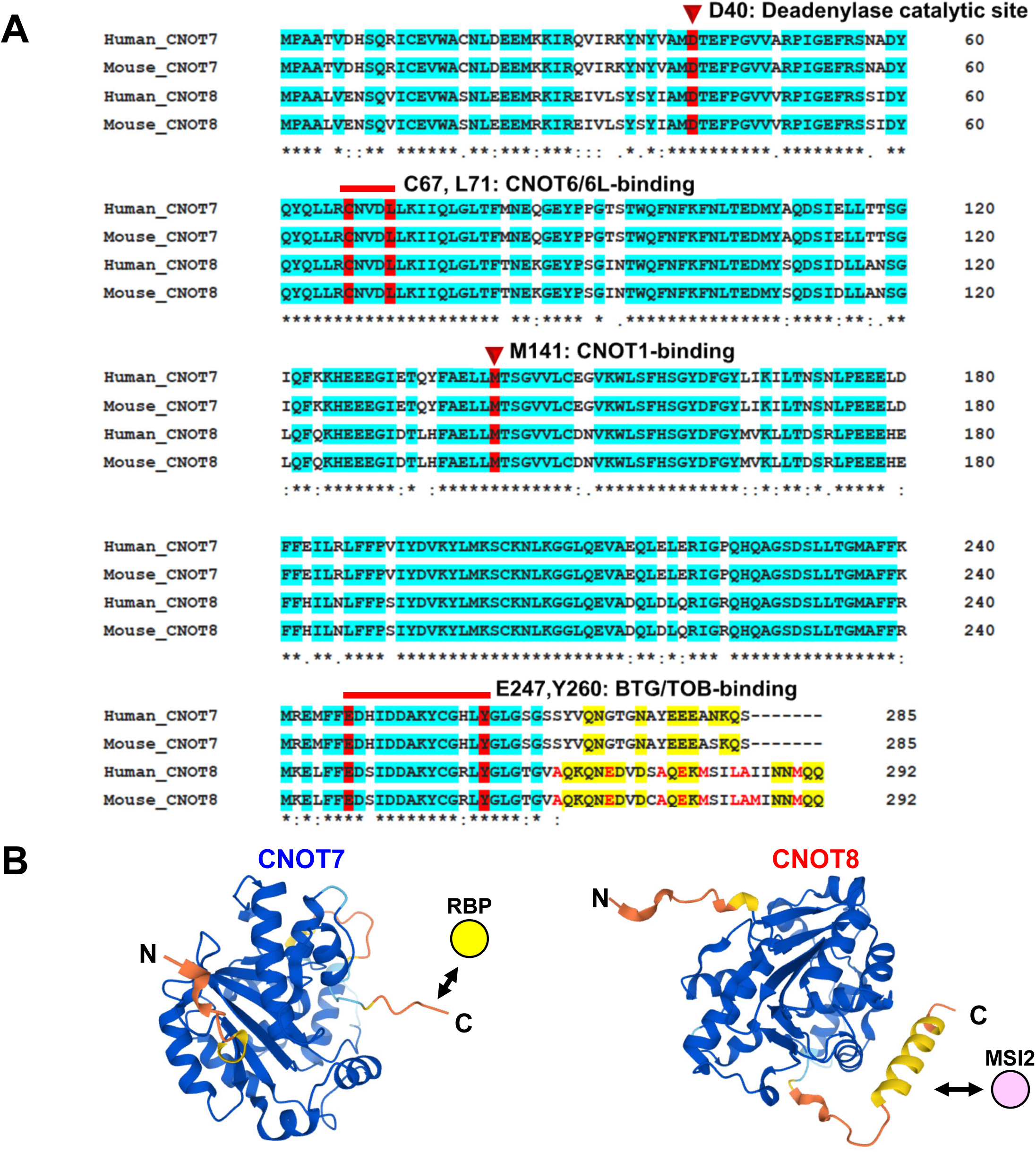
Distinctive amino acid sequences between CNOT7 and CNOT8 at their C-terminus, related to Figure 6. **(A)** Amino acid sequence alignments between human CNOT7, mouse CNOT7, human CNOT8 and mouse CNOT8. The deadenylase catalytic site (D40), the CNOT6/6L-binding site (C67, L71), the CNOT1-binding site (M141) and the BTG/TOB family protein binding site (E247, Y260) are shown in red. Positive and negative charged amino acids in the intrinsically disordered domains at the C-termini of CNOT7 and CNOT8 were highlighted in yellow. Amino acids at the C-terminus of CNOT8 that preferentially form an α-helix are indicated in red. **(B)** 3D structures of CNOT7 and CNOT8 predicted by AlphaFold2. Distinctive 3D structures of CNOT7 and CNOT8 at the C-termini, in which possible distinctive RNA-binding proteins including MSI2 interact with. An α-helix is present at the C-terminus of CNOT8, whereas that is absent at that of CNOT7.

**Figure S6.**
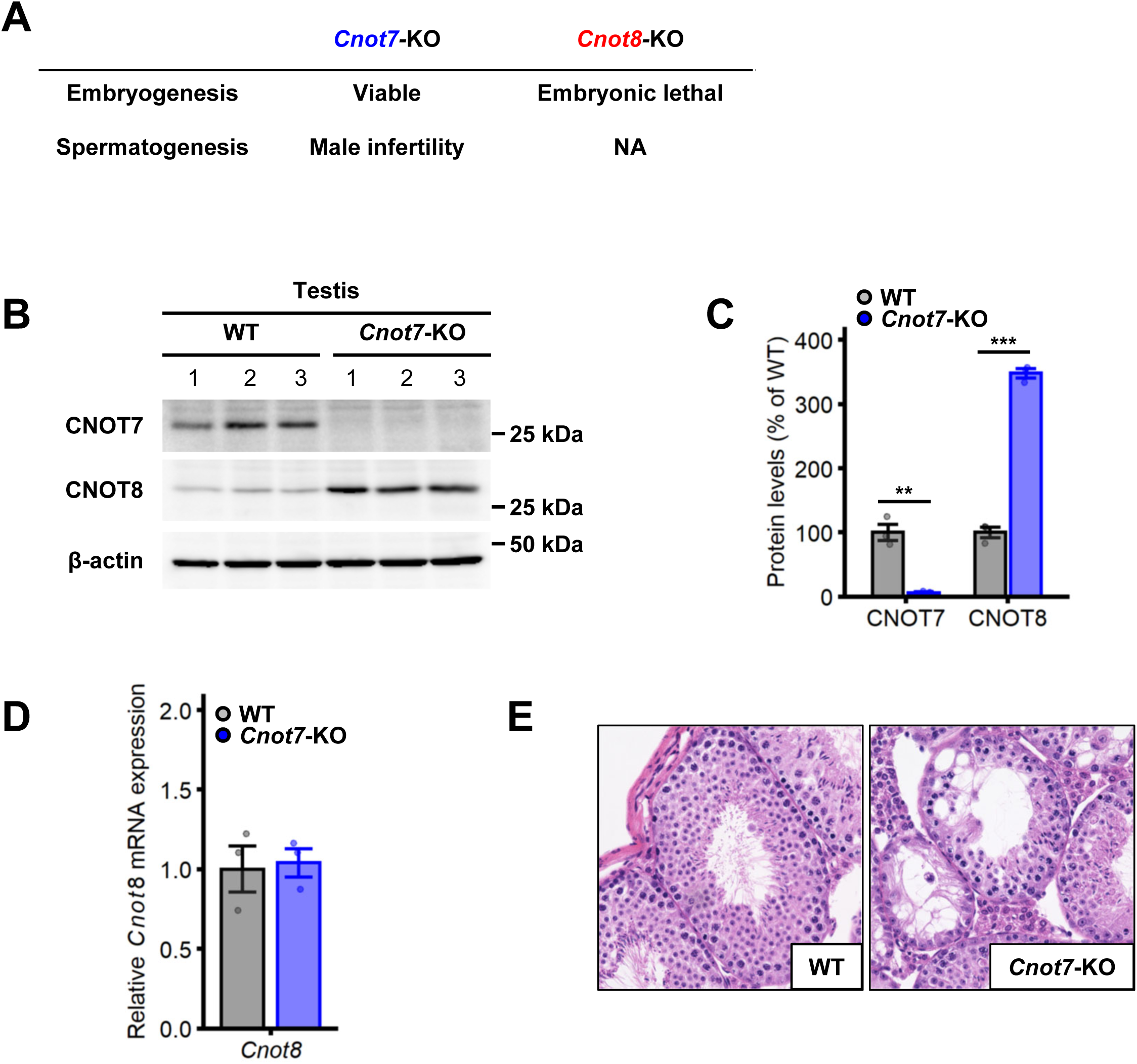
Distinctive physiological functions between CNOT7 and CNOT8, related to discussion. **(A)** Distinctive physiological functions between *Cnot7*-KO and *Cnot8*-KO mice in embryogenesis and spermatogenesis. *Cnot7*-KO mice are viable, whereas *Cnot8*-KO mice are embryonic lethal. *Cnot7*-KO mice exhibit male infertility with abnormal sperm formation. **(B)** Elevated CNOT8 expression in *Cnot7*-KO testes. CNOT7, CNOT8 and β-actin in the testes of WT (n=3) and *Cnot7*-KO mice (n=3) are shown by western blotting. **(C)** Quantitative analysis of protein bands in panel S6B. Band intensities of CNOT7 and CNOT8 in the testes of WT (grey) and *Cnot7*-KO (blue) mice were measured by NIH ImageJ. Values of the average of WT mice were set as 100 %. Data represent means ±SEM. One-Way ANOVA, post-hoc Tukey test, *p<0.05; **p<0.01; and ***p<0.001. **(D)** Relative *Cnot8* mRNA expression in WT (grey; n=3) and *Cnot7*-KO testes (blue; n=3) was determined by qPCR. The values of WT testes were set as 1. Data represent means ±SEM. **(E)** Abnormal spermatogenesis in *Cnot7*-KO testis. Histology of WT and *Cnot7*-KO testis sections were analyzed by HE staining.

